# Scalable generation of pure CD103+ cDC1 from iDC1 cultures

**DOI:** 10.64898/2026.06.04.730212

**Authors:** Sukriti Sharma, Fiona Flynn, Brian Capaldo, Ronald Holewinski, Qingrong Chen, Daoud Meerzaman, Thorkell Andresson, Christian T. Mayer

## Abstract

Conventional type 1 dendritic cells (cDC1) specialize in cross-presentation and interleukin-12 production and are critical for immunity against intracellular pathogens and tumors, but remain rare *in vivo*, limiting mechanistic and translational studies. Existing bone marrow-derived dendritic cell (BMDC) methods do not achieve highly selective enrichment of cDC1 or scalable production at high purity. Here, we established a novel *in vitro* culture system for selective generation of CD103+ cDC1 from mouse bone marrow using defined media conditions together with recombinant FLT3L, GM-CSF, and Kit ligand (KitL), termed iDC1. iDC1 cultures enabled scalable generation of an estimated 1.5 x 10^9^ CD103+ cDC1 at greater than 95% purity from a single mouse, representing at least a 75-fold increase relative to previous recombinant cytokine-based methods. Phenotypic and transcriptional analyses demonstrated that iDC1 closely align with the CD103+ cDC1 lineage while remaining clearly distinct from macrophage populations. Functionally, iDC1 responded robustly to innate stimulation, produced interleukin-12 and inflammatory chemokines, and efficiently cross-presented cell-associated antigen to CD8+ T cells. Mechanistically, KitL and GM-CSF regulated distinct stages of cDC1 generation, whereas proteomic, phospho-proteomic, and functional analyses demonstrated that GM-CSF suppresses apoptosis and oxidative stress while promoting cDC1 proliferation. iDC1 generation was dependent on the +32 kb *Irf8* enhancer required for *bona fide* cDC1 development, and STAT5-and BRD4-associated regulatory programs were identified as important regulators of efficient iDC1 generation. Together, these findings establish iDC1 cultures as a scalable platform for studying cDC1 biology and developing cDC1-based immunotherapeutic strategies.

## Introduction

Dendritic cells (DC) comprise conventional DCs (cDC) and plasmacytoid DCs (pDC). Two major cDC subsets, cDC1 and cDC2, populate both lymphoid and non-lymphoid tissues and can be broadly classified as resident or migratory cDC, respectively^1^. cDC1 specialize in the cross-presentation of cell-associated antigens, produce interleukin-12 following toll-like receptor (TLR) 3, TLR11 and TLR12 activation, and orchestrate adaptive immune responses to intracellular pathogens and tumors^2–5^. Significant progress has been made in defining the transcriptional network controlling cDC1 lineage specification. In particular, transcription factor IRF8 is essential for cDC1 lineage development through formation of complexes with basic leucine zipper transcription factors BATF, BATF2 or BATF3 together with Jun proteins, resulting in IRF8 autoactivation at the +32 kb *Irf8* enhancer^6,7^. Mice lacking the +32 kb *Irf8* enhancer (Irf8+32^-/-^) completely lack cDC1 under both steady-state and inflammatory conditions^8^.

Because DCs, and particularly cDC1, are rare *in vivo*, bone marrow-derived DC (BMDC) culture systems have become invaluable tools for studying DC development, biology, and therapeutic potential. Culture of bone marrow cells with GM-CSF generates macrophages (GM-Mac) and DC-like cells (GM-DC) that do not clearly correspond to cDC1 or cDC2 populations found *in vivo*^9–11^. In contrast, FLT3L-driven cultures (FLT3L-DC) generate all major DC subsets, including pDC, cDC1 and cDC2^12,13^.

Recent efforts have focused on selectively biasing BMDC differentiation toward cDC1^14–17^, which could greatly facilitate basic and pre-clinical research by increasing cDC1 yield and purity while potentially circumventing the need for magnetic enrichment or fluorescence-activated cell sorting. Low-dose GM-CSF combined with FLT3L strongly biases iCD103-DC cultures toward cDC1 production^16^. Kit ligand/stem cell factor (KitL), interleukin-4 (IL-4), and OP9 stromal cells have also been reported to enhance FLT3L-dependent cDC1 generation^14,15,17^. However, currently available methods do not achieve highly selective cDC1 production above 95% purity at scalable yield. Importantly, BMDC purity should not be assessed solely within gated cDC populations, but must also account for macrophage and other non-DC contaminants contributing to overall culture heterogeneity.

Here, we describe a 23-day BMDC method termed iDC1 that uses recombinant cytokines to generate approximately 1.5 x 10^9^ CD103+ cDC1 at greater than 95% purity from a single mouse. We further show that iDC1 are +32 kb *Irf8* enhancer-dependent, exhibit phenotypic, transcriptional, and functional properties of *bona fide* cDC1 and identify GM-CSF-regulated signaling and regulatory programs that promote efficient cDC1 generation and maintenance. These findings establish iDC1 cultures as a scalable platform for studying cDC1 biology and therapeutic potential.

## Methods

### Mice

C57BL/6J (Strain #000664), C57BL/6-Irf8em1Kmm/J (Irf8+32^-/-^; Strain #032744)^8^, B6N(129S4)-Xcr1tm1.1(cre)Kmm/J (Xcr1^Cre^; Strain #035435)^18^, and B6.C-Tg(CMV-cre)1Cgn/J (CMV^Cre^; Strain #006054)^19^ mice were purchased from The Jackson Laboratory. Rosa26^LSL-Bcl2-IRES-GFP^ mice^20^ were provided by Dr. Hamid Kashkar (University of Cologne). Rosa26^INDIA^ mice^21^ were provided by Dr. Michel Nussenzweig (The Rockefeller University). B6.129S6-Stat5btm1Mam Stat5atm2Mam/Mmjax (STAT5a/b^fl/fl^) mice^22^ were provided by Dr. Hyun Park (National Cancer Institute). Brd4^fl/fl^ mice^23^ were provided by Dr. Dinah Singer (National Cancer Institute). C57BL/6-Tg(TcraTcrb)1100Mjb/J (OT-I) mice^24^ expressing CD45.1 were provided by Dr. Alfred Singer (National Cancer Institute). Rosa26^LSL-Fucci2aR^ mice^25^ were provided by Dr. Ettore Appella (National Cancer Institute) and crossed with CMV^Cre^ mice to induce germline deletion of the loxP-STOP-loxP cassette. Following crossing to C57BL/6J mice, Cre-negative Rosa26^Fucci2aR^ mice exhibiting ubiquitous Fucci2aR expression were subsequently intercrossed to homozygosity. Mice were maintained under specific pathogen-free conditions at NCI (Frederick and Bethesda) on a 12 h dark/light cycle. Male and female mice aged 6–18 weeks were used for all experiments.

### Production and purification of recombinant cytokines

A pcDNA3-based expression plasmid containing a human Fc tag (Addgene plasmid #141183) was modified to generate expression vectors encoding recombinant FLT3L or GM-CSF. The FLT3L construct encoded a murine immunoglobulin signal peptide^26^, an Avi-tag, an N-terminal human Fc tag, and human FLT3L (Accession number AAA17999.1) amino acids 27 to 185. The GM-CSF construct encoded a murine immunoglobulin signal peptide^26^, murine GM-CSF (Accession number Q14AD9) amino acids 18 to 141, a C-terminal human Fc tag, and an Avi-tag. Recombinant cytokines were produced by transient transfection of 293F cells and purified using Protein G Sepharose 4 Fast Flow (GE Healthcare) as previously described^26,27^. Purified proteins were buffer exchanged into PBS, sterile filtered, and analyzed by SDS-PAGE to confirm protein integrity. Protein concentrations were measured with NanoDrop One (Thermo Fisher).

### Generation of bone marrow-derived dendritic cells (BMDCs) and induced CD103+ cDC1 (iDC1)

Unless otherwise indicated, complete culture medium consisted of RPMI 1640 containing GlutaMAX or L-glutamine (Gibco #61870036 or Fisher Scientific #FB12999107) supplemented with 10% heat-inactivated fetal bovine serum (FBS; Biofluids #200P-500, lot #814037 or GeminiBio #S11150, lot #K23233), 1x Penicillin-Streptomycin (Gibco #15070063), 1x 2-mercaptoethanol (Gibco #21985023), 25 mM HEPES (Quality Biological #118-089-721 or Gibco #15630080), 2 mM additional GlutaMAX (Gibco #35050061), and 1 mM sodium pyruvate (Gibco #11360070). FBS lots were pre-screened for efficient BMDC generation. Where indicated, complete medium consisted of RPMI 1640, DMEM containing high glucose and GlutaMAX (Gibco #10569044), or IMDM (Gibco #12440053), each supplemented with 10% heat-inactivated FBS, 1x Penicillin-Streptomycin, and 1x 2-mercaptoethanol. Media were additionally supplemented where indicated with 1x MEM non-essential amino acids (Gibco #11140050) and 1x MEM amino acids (Gibco #11130051).

Femurs and tibiae were harvested from mice of the indicated genotypes, cut at both ends, and bone marrow was flushed with complete RPMI 1640 medium. Cells were passed through a 70 μm cell strainer, centrifuged for 7 min at 1300 rpm, and subjected to red blood cell lysis using ACK lysis buffer (Quality Biological #118-056-101 or KD Medical #RGF-3015). Cells were subsequently washed and resuspended in complete RPMI 1640 medium.

GM-CSF-derived DCs (GM-Mac/DC), FLT3L-derived DC (FLT3L-DC), iCD103-DC and FLT3L/KitL-derived DC were generated as described previously^10,11,13,16,17^, except that the optimized culture media described above and recombinant GM-CSF and FLT3L produced in-house were used. All BMDC cultures were established in 100 mm x 15 mm non-tissue-culture-treated bacteriological Petri dishes (Falcon #351029 or Kord-Valmark #KORD-2910P) and maintained at 37 °C and 5% CO.

For GM-Mac/DC cultures, 2 x 10^6^ bone marrow cells were cultured in 12 ml complete RPMI medium containing 20 ng/ml GM-CSF. On day 2, 10 ml fresh complete medium was added. On days 4, 6, and 8, 10 ml culture supernatant was collected, centrifuged, and cell pellets were resuspended in 10 ml fresh complete medium containing 20 ng/ml GM-CSF and returned to the original plates. Non-adherent cells were harvested on day 9.

For FLT3L-DC cultures, 15 x 10^6^ cells were cultured in 12 ml complete RPMI medium containing 200 ng/ml FLT3L for 9 days.

For KitL/FLT3L-DC cultures, 7.5 x 10^6^ cells were cultured in 12 ml complete RPMI medium containing 200 ng/ml FLT3L and 200 ng/ml recombinant murine KitL (PeproTech #250-03). On day 3, cells were harvested, centrifuged, resuspended in fresh medium containing 200 ng/ml FLT3L, and returned to the original plates. Cells were harvested on day 11.

For iCD103-DC cultures, 15 x 10^6^ cells were cultured in 12 ml complete RPMI medium containing 200 ng/ml FLT3L and 2 ng/ml GM-CSF. Non-adherent cells were harvested on day 9, counted, centrifuged, and 3 x 10^6^ cells were replated in 12 ml complete RPMI medium containing 200 ng/ml FLT3L and 2 ng/ml GM-CSF. Cells were harvested on day 16.

For iDC1 cultures, 15 x 10^6^ cells were cultured in complete RPMI medium containing 200 ng/ml FLT3L, 2 ng/ml GM-CSF, and 200 ng/ml KitL (PeproTech #250-03) for 9 days. Non-adherent cells were harvested on day 9, counted, centrifuged, and 3 x 10^6^ cells were replated in 12 ml complete RPMI medium containing 200 ng/ml FLT3L and 2 ng/ml GM-CSF. The day 9 replating procedure was repeated on day 16, and non-adherent iDC1 were routinely harvested on day 23. For selected experiments, iDC1 cultures were established in 150 mm x 15 mm non-tissue-culture-treated bacteriological Petri dishes (Falcon #351058) in parallel, in which case culture volume and cell numbers were scaled proportionally to surface area (2.6-fold).

### Light microscopy

Light microscopy images were acquired directly from iDC1 culture dishes using an Axio Vert.A1 inverted microscope (Zeiss) equipped with 10x or 20x objectives and a Canon EOS Rebel SL3 camera.

### Flow cytometry

For details about the antibodies used see Supplementary Table S1. Flow cytometry data were acquired on a FACSymphony A5 (BD Biosciences) and analyzed using FlowJo version 10 (BD Biosciences). For live-cell staining, Fc receptors were blocked for 15 min on ice using anti-CD16/CD32 antibody (2.4G2, produced in-house). Cells were centrifuged for 3 min at 1300 rpm and stained with antibodies against surface antigens for 45 min at 4 °C, followed by three washes with FACS buffer (PBS containing 0.1% w/v BSA and 2 mM EDTA). Where indicated, cells were subsequently stained with APC-Fire750-conjugated streptavidin (BioLegend #405250; 1:320 dilution) for 10 min at 4 °C and washed three times. Cells were resuspended in FACS buffer containing 0.2 μg/ml propidium iodide (PI; Sigma-Aldrich #P4170) or 0.1 μg/ml DAPI (Sigma-Aldrich #D9542) before acquisition to exclude dead cells. For selected experiments, cells were incubated with 5 μM CellROX (Thermo Fisher Scientific #C10422) for 30 min at 37 °C, washed three times with PBS, and subsequently stained for surface markers.

For intracellular staining of EdU, active caspase-3 (aCasp3), and BRD4, cells were washed twice with PBS and stained with Zombie UV Fixable Viability Dye (BioLegend #423108; 1:500 dilution in PBS) for 30 min at 4 °C before Fc receptor blocking and surface staining.

For detection of EdU and aCasp3, cells were fixed and permeabilized using BD Cytofix/Cytoperm (BD #554714) for 30 min at 4 °C, followed by two washes with 1x Perm/Wash buffer (BD #554714). EdU incorporation was detected using Click-iT Plus EdU Alexa Fluor 488 Flow Cytometry Assay Kit (Thermo Fisher #C10632) according to the manufacturer’s instructions. Cells were incubated with Click reaction mix for 30 min in the dark and washed twice with 1x Perm/Wash buffer. Where indicated, cells were subsequently stained with anti-aCasp3 antibody diluted in 1x Perm/Wash buffer for 45 min at 4 °C, washed twice with 1x Perm/Wash buffer, and resuspended in FACS buffer.

For detection of BRD4, cells were fixed with 2% paraformaldehyde (PFA; Electron Microscopy Sciences #15714-S) for 10 min at room temperature, washed twice with PBS, and permeabilized with pre-chilled methanol (Thermo Fisher #A412-1) on ice for 20 min. After two washes with FACS buffer, cells were incubated with rabbit anti-BRD4 antibody (Abcam #ab289886) for 30 min at room temperature followed by two washes with FACS buffer. Cells were then incubated with goat anti-rabbit Alexa Fluor 488 antibody (Abcam #ab150081; 1:1000 dilution) for 30 min at room temperature, washed twice with FACS buffer, and resuspended in FACS buffer.

For detection of phosphorylated STAT5 (p-STAT5), cells were resuspended in sterile RPMI 1640 medium and rested for 30 min at 37 °C before stimulation with 2 ng/ml GM-CSF for 30 min at 37 °C. Cells were fixed with 1.6% PFA for 10 min at 37 °C, washed twice with HBSS (Gibco #14170-112) supplemented with 10% heat-inactivated FBS (Gemini #S11150, lot #K23233), and permeabilized with pre-chilled methanol (Thermo Fisher #A412-1) for 15 min on ice. After two washes with HBSS containing 10% FBS, cells were incubated with Fc block and p-STAT5 antibody for 30 min at room temperature followed by staining with antibodies against surface antigens for 20 min at room temperature. Cells were subsequently stained with APC-Fire750-conjugated streptavidin (BioLegend #405250; 1:320 dilution) for 10 min at room temperature, washed twice with FACS buffer, and resuspended in FACS buffer.

### Proteomics

iDC1 were generated from Xcr1^Cre^Rosa26^LSL-Bcl2-IRES-GFP^ bone marrow to prevent apoptosis following GM-CSF withdrawal (Supplementary Figure S3), harvested on day 23, and washed twice with complete RPMI 1640 medium. For each biological replicate, multiple plates each containing 3 x 10^6^ iDC1 in 12 ml complete RPMI 1640 medium and either 200 ng/ml FLT3L alone or 200 ng/ml FLT3L plus 2 ng/ml GM-CSF were prepared and cultured for an additional 20 h at 37 °C and 5% CO. Non-adherent iDC1 were then harvested, pooled, washed twice with PBS, and cell pellets from seven biological replicates per condition were frozen prior to proteomic analysis.

Each cell pellet was lysed in 500 μl EasyPep Lysis Buffer (Thermo Fisher #A45735) supplemented with 1x phosphatase inhibitor (Thermo Fisher #A32957) and universal nuclease (Thermo Fisher #88700). Protein concentration was determined by BCA assay and 150 μg protein from each sample was used for digestion. Two control and two GM-CSF-treated samples were digested twice to complete the TMTpro plex and increase material available for phospho-peptide enrichment. Samples were adjusted to 150 μl total volume with lysis buffer and mixed with 50 μl digestion buffer containing 50 mM TCEP, 200 mM chloroacetamide, and 58 ng/μl trypsin/LysC in 75 mM HEPES (pH 8.0). Samples were incubated at 37 °C for 21 h and subsequently labeled with TMTpro 18-plex reagents (Thermo Fisher #A52045; lots 3214388 and 3214389) for 1 h at 25 °C. Excess TMTpro reagent was quenched with 5% hydroxylamine and 20% formic acid for 10 min, after which samples were combined. Peptides were cleaned using EasyPep Maxi columns (Thermo Fisher #A45734) according to the manufacturer’s instructions and eluted in 3 ml elution buffer. Eluted peptides were divided into a 50 μl aliquot for global proteomic analysis and four 737 μl aliquots for phosphopeptide enrichment and subsequently dried.

Phospho-peptides were sequentially enriched using High-Select TiO (Thermo Fisher #A32993) and High-Select Fe-NTA IMAC (Thermo Fisher #A32992) kits according to the manufacturer’s instructions. Global and phospho-enriched peptide samples were resuspended in 0.1% formic acid and analyzed in duplicate using a Dionex U3000 RSLC coupled to an Orbitrap Eclipse mass spectrometer (Thermo Fisher Scientific) equipped with an EasySpray ion source and FAIMS Duo interface. Peptides were separated by reverse-phase liquid chromatography and analyzed using Orbitrap-based data-dependent acquisition with TurboTMT quantification.

All MS files were processed in Proteome Discoverer 2.4 using the Sequest search engine and searched against the UniProt mouse database. Searches were performed using full tryptic specificity with up to two missed cleavages, a precursor mass tolerance of 10 ppm, and a fragment mass tolerance of 0.02 Da. Variable modifications included methionine oxidation (+15.995 Da) and serine, threonine, and tyrosine phosphorylation (+79.966 Da), whereas carbamidomethylation of cysteine (+57.021 Da) and TMTpro labeling of lysine residues and peptide N-termini (+304.207 Da) were specified as fixed modifications. Percolator was used for false discovery rate (FDR) analysis. Phosphosite localization was performed using IMP-ptmRS with a localization threshold of 90%. TMTpro reporter ions were quantified using the Reporter Ion Quantifier node and normalized on total peptide intensity for each channel.

Statistical analyses were performed in Proteome Discoverer 2.4 using Log -transformed median intensities and ANOVA-based p-value calculation. Differentially abundant proteins and phospho-peptides were identified using cutoffs of absolute Log fold change (Log FC) > 0.6 and p < 0.05. Proteins or peptides detected in at least two replicates of one condition and absent from all replicates of the comparison condition were assigned adjusted Log FC and p-values to permit inclusion in downstream analyses. TMTpro channel assignments are provided in Supplementary Data S5.

GM-CSF-regulated differentially abundant proteins were analyzed using Ingenuity Pathway Analysis, and enriched transcription factor target signatures were analyzed using the Database for Annotation, Visualization, and Integrated Discovery (DAVID; https://davidbioinformatics.nih.gov/).

### FACS sorting and RNA sequencing analysis

BMDCs were generated as described above and either left unstimulated or stimulated overnight (∼16 h) with 10 µg/ml Poly(I:C) (InvivoGen #tlrl-pic). Non-adherent BMDCs were harvested, stained for flow cytometry as described above, and directly sorted into Trizol LS Reagent (Thermo Fisher #10296010) using a FACSymphony S6 (BD Biosciences).

Three to four biological replicates of each population were sorted after excluding DAPI^+^ cells. GM-Mac/DC cultures were gated as CD11c+CD11b+I-A/I-E(low)CD64+CD115+ macrophages. FLT3L, iCD103-DC and iDC1 cultures were gated as CD64-CD11c+CD45R/B220-CD103+CD172a-Xcr1+CD24+ cDC1. Additional populations sorted from FLT3L-DC cultures included CD64-CD11c+CD45R/B220-CD103-CD172a-Xcr1+CD24+ cDC1, CD64-CD11c+CD45R/B220-CD103-CD172a+Xcr1-CD24(low) cDC2, and CD64+CD172a+ macrophages.

For tissue DC and macrophage isolation, spleens and lungs from C57BL/6 mice were injected with digestion buffer containing HBSS (Gibco #14170-112), 4 mM calcium chloride (Quality Biological #351-130-721), 1 mM magnesium chloride (Quality Biological #351-033-721), and 10% heat-inactivated FBS (Biofluids #200P-500, lot #814037). Spleens were digested with 1 mg/ml Collagenase D (Millipore Sigma #11088858001) and 0.1 mg/ml DNase I (Millipore Sigma #11284932001), whereas lungs were digested with 0.2 mg/ml Collagenase IV (Millipore Sigma #C4-22-1g) and 0.1 mg/ml DNase I. Tissues were minced and incubated at 37 °C for 25 min (spleen) or 90 min (lung). Subsequently, 10 mM EDTA (Invitrogen #15575-038) was added for 5 min and tissues were passed through a 70 μm cell strainer.

Single-cell suspensions were centrifuged for 7 min at 1300 rpm and resuspended in high-density medium consisting of three parts HBSS and one part OptiPrep (Serumwerk Bernburg AG #1893). Suspensions were overlaid with low-density medium consisting of one part OptiPrep and 4.2 parts FACS buffer, followed by a final HBSS overlay. Density gradients were centrifuged for 15 min at 1500 rpm and 4 °C without brake. Low-density cells enriched for DCs and macrophages were collected from the interphase, washed with FACS buffer, stained for flow cytometry as described above, and directly sorted into TRIzol LS Reagent using a FACSymphony S6.

Lineage markers included CD19, Ter-119, and NK1.1. Three biological replicates of each population were sorted after exclusion of DAPI^+^ cells. Splenic populations included Lineage-CD45R/B220-I-A/I-E(high)CD11c(high)CD3ε-CD45+CD11b-Xcr1+ cDC1, Lineage-CD45R/B220-I-A/I-E(high)CD11c(high)CD3ε-CD45+CD11b+Xcr1- cDC2, and Lineage-CD45R/B220-CD11c(low)I-A/I-E(low)F4/80+CD11b+CD3ε-CD45+ red pulp macrophages. Lung alveolar macrophages were gated as Lineage-CD45+SiglecF+CD11c+ cells.

Total RNA was extracted according to the manufacturer’s instructions for Trizol-LS. 1µl Pellet Paint (Novagen #69049) was added to the aqueous phase prior to RNA precipitation with isopropanol to facilitate precipitation and pellet visualization. Extracted RNA was further purified with the Nucleospin RNA XS kit (Takara Bio #740902.50) according to the manufacturer’s instructions. Subsequently, 100 pg total RNA was processed using the SMARTer Ultra Low Input RNA Kit (Takara) to generate cDNA libraries. Sequencing libraries were then prepared using the Nextera XT DNA Sample Preparation Kit (Illumina). Sixty samples were pooled and sequenced on a NovaSeq X Plus 1.5B instrument using paired-end sequencing.

Raw fastq files were aligned and quantified using the NF Core RNA seq pipeline (https://zenodo.org/records/20072251). Quantified reads were analyzed by using tximport^28^ to import reads and annotations, and counts normalized using edgeR^29,30^. Limma-voom^31^ and variancePartition^32,33^ were used to perform linear modeling with mixed effects to identify differentially expressed genes between conditions. Gene set enrichment analysis was performed using clusterProfiler^34^ with gene sets obtained using msigdbr (https://CRAN.R-project.org/package=msigdbr). ImmGen database was accessed using celldex and compared against limma-voom transformed data using SingleR^35^.

### iDC1 stimulation

Non-adherent day 23 iDC1 were harvested, counted, washed once with sterile PBS, and resuspended in complete RPMI 1640 medium. Subsequently, 2 x 10^5^ iDC1 were plated in 200 μl complete RPMI 1640 medium in non-tissue-culture-treated 96-well round-bottom plates and stimulated with 100 ng/ml LPS (Cat. L2630, Sigma), 10 μg/ml poly(I:C) (Cat. tlrl-pic, InvivoGen), 1 μM CpG-ODN 1826 (Cat. tlrl-1826, InvivoGen), combined poly(I:C) plus CpG-ODN 1826, or left untreated for 21–22 h at 37 °C and 5% CO. Cell pellets were analyzed for co-stimulatory marker expression by flow cytometry. Culture supernatants were analyzed using the Proteome Profiler Mouse XL Cytokine Array Kit (Cat. ARY028, R&D Systems) according to the manufacturer’s instructions. Array membranes were developed using SuperSignal West Pico PLUS Chemiluminescent Substrate (Thermo Fisher #34577), and chemiluminescent signals were imaged using a UVP ChemStudio system (Analytik Jena). Images were analyzed using ImageJ. Relative protein expression levels were quantified by measuring spot mean gray values, which were averaged across duplicate spots and two independent experiments. Spots with average mean gray values greater than 20 were considered positive.

### Preparation of heat-killed Listeria monocytogenes expressing OVA

All agar plates and liquid media were supplemented with 50 μg/ml erythromycin (Sigma #E5389). Frozen *Listeria monocytogenes* expressing OVA (LM-OVA)^36^ were streaked onto BHI agar plates and incubated overnight at 37 °C. A single colony was inoculated into 4 ml BHI medium and cultured overnight at 37 °C and 225 rpm. The following day, 150 ml pre-warmed BHI medium was inoculated to an OD_600_ of approximately 0.1 and cultured at 37 °C and 225 rpm until reaching an OD_600_ of approximately 0.5–1.5. Cultures were centrifuged for 20 min at 4000 rpm, and bacterial pellets were washed twice with 50 ml sterile PBS by centrifugation for 10 min at 4000 rpm. LM-OVA were resuspended to an estimated concentration of 1 x 10^11^ CFU/ml based on the approximation that an OD_600_ of 0.1 corresponds to 2 x 10^8^ CFU/ml^37^. LM-OVA stocks were heat-inactivated by incubation for 1 h at 70 °C (HKLM-OVA) and stored at −80 °C. Complete inactivation was confirmed by absence of colony growth on BHI agar plates.

### In vitro antigen cross-presentation assay

Non-adherent day 23 WT and Irf8+32^-/-^ iDC1 were harvested, counted, and resuspended in complete RPMI 1640 medium. B cells were purified from C57BL/6 spleens following red blood cell lysis with ACK lysis buffer (Quality Biological #118-056-101) using Mouse CD43 Microbeads (Miltenyi Biotec #130-049-801) and LS columns (Miltenyi Biotec #130-042-401) according to the manufacturer’s instructions. Purified B cells were counted and resuspended in complete RPMI 1640 medium. CD8+ T cells were purified from spleens and lymph nodes of CD45.1+ OT-I mice using the Mouse CD8a+ T cell Isolation Kit (Miltenyi Biotec #130-104-075) and LS columns according to the manufacturer’s instructions. Purified CD8+ T cells were washed with sterile PBS and labeled with 5 µM CellTrace Violet (CTV; Invitrogen #C34557) for 20 min at 37 °C according to the manufacturer’s instructions. Cells were subsequently counted, centrifuged, and resuspended in complete RPMI 1640 medium. For cross-presentation assays, 2.5 x 10^4^ CTV-labeled OT-I T cells were cultured with 1 x 10^4^ iDC1 or 1 x 10^4^ B cells in the presence of either 10^8^ Heat-killed *Listeria monocytogenes* expressing OVA^36^ (HKLM-OVA), 10 µg/ml soluble OVA (Sigma #A5503), 1 µg/ml SIINFEKL peptide (OVA_257-264_; Life Technologies), or no stimulation. After 4 days at 37 °C, proliferation of CTV(low)CD44+CD8α+CD45.1+Vα2+ OT-I T cells was analyzed by flow cytometry as percentages and total cell numbers.

### Statistical analysis

Statistical analyses were performed by GraphPad Prism software (version 10). Data were assessed for normality using the Anderson-Darling, D’Agostino-Pearson, Shapiro-Wilk, and Kolmogorov-Smirnov tests where applicable. Unpaired two-tailed Student’s *t*-tests were used for normally distributed data or when sample size was insufficient for normality testing. No adjustments were made for multiple comparisons. Statistical tests used are indicated in the figures and figure legends. Experiments were performed independently two to three times.

## Results and Discussion

### Culture media type and composition regulate cDC1 production

We first assessed whether culture conditions influence cDC1 production using previously described iCD103-DC cultures generated with FLT3L and low-dose GM-CSF that strongly enrich for CD103+ cDC1^16^. RPMI 1640 generated significantly more cDC1 than IMDM and DMEM (Supplementary Figure S1A-C), indicating that basal media composition strongly influences cDC1 differentiation efficiency. Compared with RPMI 1640, IMDM and DMEM contain higher concentrations of amino acids. Consistently, supplementation of RPMI 1640 with excess essential and non-essential amino acids significantly reduced cDC1 generation relative to control RPMI 1640 cultures (Supplementary Figure S1D–F), suggesting that elevated amino acid availability suppresses efficient cDC1 differentiation. These findings suggest that cDC1 differentiation is highly sensitive to extracellular metabolic conditions. Elevated amino acid availability may alter nutrient-sensing and metabolic pathways required for efficient cDC1 differentiation, including pathways linked to mTOR signaling^38^ and cellular biosynthetic activity.

In contrast, supplementation of RPMI 1640 with HEPES, sodium pyruvate, and the stable dipeptide L-glutamine substitute GlutaMAX (this supplement mixture is abbreviated as HSpG) significantly increased cDC1 production (Supplementary Figure S1A–C), consistent with metabolic support and buffering capacity promoting cDC1 expansion and/or survival. Based on these findings, RPMI 1640 supplemented with HSpG was used for subsequent experiments. Together, these findings identify extracellular metabolic conditions as an important determinant of efficient cDC1 generation and establish RPMI 1640 supplemented with HSpG as an optimized basal medium for subsequent iDC1 cultures.

### iDC1 cultures produce large numbers of +32 kb Irf8 enhancer-dependent cDC1 of high purity

We next established a distinct BMDC culture system incorporating the defined RPMI 1640 medium supplemented with HSpG together with FLT3L, low-dose GM-CSF, and KitL during the first 9 days of culture, followed by serial replating with FLT3L and GM-CSF every 7 days (termed iDC1; Figure 1A). For comparison, previously described iCD103-DC cultures generated with FLT3L and low-dose GM-CSF that effectively enrich for CD103+ cDC1^16^ were analyzed in parallel using the same medium. Flow cytometric analysis at days 16, 23, and 30 showed that iDC1 cultures exhibited significantly higher cDC1 purity and markedly greater cDC1 yields compared with iCD103-DC cultures at all time points (Figure 1B, C). Overall cDC1 purity increased from ∼60% in day 16 iCD103-DC cultures to ∼97% in day 23 iDC1 cultures (Figure 1B), with total cDC1 yields peaking at day 23 (Figure 1C), representing an approximately 75-fold increase relative to the original iCD103-DC method using recombinant cytokines^16^. Day 23 was therefore selected for subsequent experiments. iDC1 cultures established in 100 mm dishes could be scaled to 150 mm dishes with only a modest reduction in yield and purity, while remaining highly efficient for cDC1 generation (Supplementary Figure S1G-I). CD103+CD172a- iDC1 expressed canonical cDC1 markers including CD24, Clec9a, and Xcr1, which were absent or expressed at lower levels in CD103-CD172a+ cDC2 present at trace amounts within the same cultures (Figure 1D, E).

**Figure 1.**
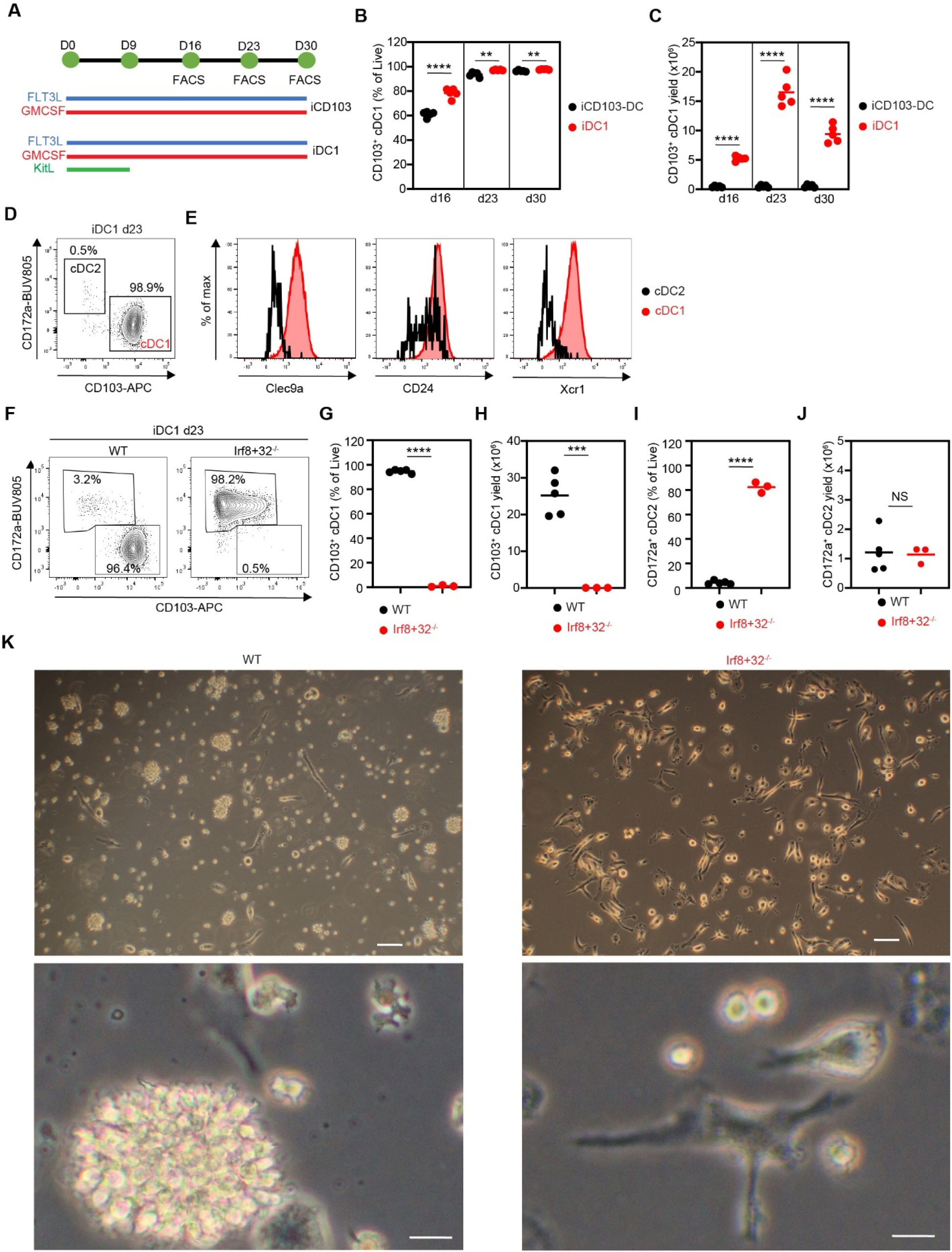
Efficient generation of pure CD103+ cDC1 in iDC1 cultures requires the +32 kb Irf8\ enhancer. (A–E) Bone marrow cells were cultured with FLT3L and GM-CSF (iCD103-DC) or with FLT3L, GM-CSF, and KitL (iDC1), followed by serial replating with FLT3L and GM-CSF every 7 days. Cells were analyzed by flow cytometry at the indicated time points. (A) Experimental scheme. (B) Frequencies and (C) total numbers of CD103+ cDC1 at the indicated time points. (D) Representative contour plot of CD103 versus CD172a expression among CD11c+CD45R/B220-CD64- cDC from day 23 iDC1 cultures. (E) Expression of indicated markers in cDC1 and cDC2 from day 23 iDC1 cultures. (F–K) iDC1 cultures were generated from WT or Irf8+32-/- bone marrow and analyzed on day 23. (F) Representative contour plots showing cDC1 and cDC2 populations among CD11c+CD45R/B220-CD64-cDC. (G) Frequency and (H) total number of cDC1. (I) Frequency and (J) total number of cDC2. (K) Representative micrographs (scale bars, 100µm (top) and 20µm (bottom)). ** p<0.01, *** p<0.001, **** p<0.0001, NS = not statistically significant (unpaired two-tailed Student’s t-test).

To determine lineage identity, iDC1 cultures were established from control and Irf8+32^-/-^ bone marrow. CD103+CD172a- cDC1 were absent in Irf8+32^-/-^ cultures compared with WT controls (Figure 1F-H), whereas remaining cells were predominantly cDC2 (Figure 1F, I) with total numbers comparable to WT controls (Figure 1J). iDC1 cultures formed floating aggregates and single cells with dendritic morphology (Figure 1K), similar to iCD103-DC cultures^16^, whereas Irf8+32^-/-^ cultures lacked floating clusters. Together, these findings establish iDC1 cultures as a scalable system for high-yield, high-purity generation of bona fide +32 kb *Irf8* enhancer–dependent cDC1.

### Comparison of iDC1 to other BMDCs, tissue DCs and macrophages

We next compared the overall cellular composition of different BMDC systems at their established endpoint culture times using a common flow cytometry gating strategy (Supplementary Figure S2A). GM-CSF-derived cultures (GM-Mac/DC) and FLT3L-derived cultures (FLT3L-DC) were analyzed on day 9, FLT3L/KitL cultures on day 11, iCD103-DC cultures on day 16, and iDC1 cultures on day 23. GM-Mac/DC cultures contained predominantly CD172a+CD64+ cells (86%) expressing high levels of CD11b and F4/80 and low levels of I-A/I-E (Figure 2A and Supplementary Figure S2), consistent with a macrophage phenotype^9^. The remaining cells could not be clearly separated from macrophages and included CD11c(high)B220- populations, most of which displayed a cDC2-like phenotype. Together, these findings confirm that GM-CSF cultures predominantly generate macrophage-like populations rather than bona fide cDC1^9,16^.

**Figure 2.**
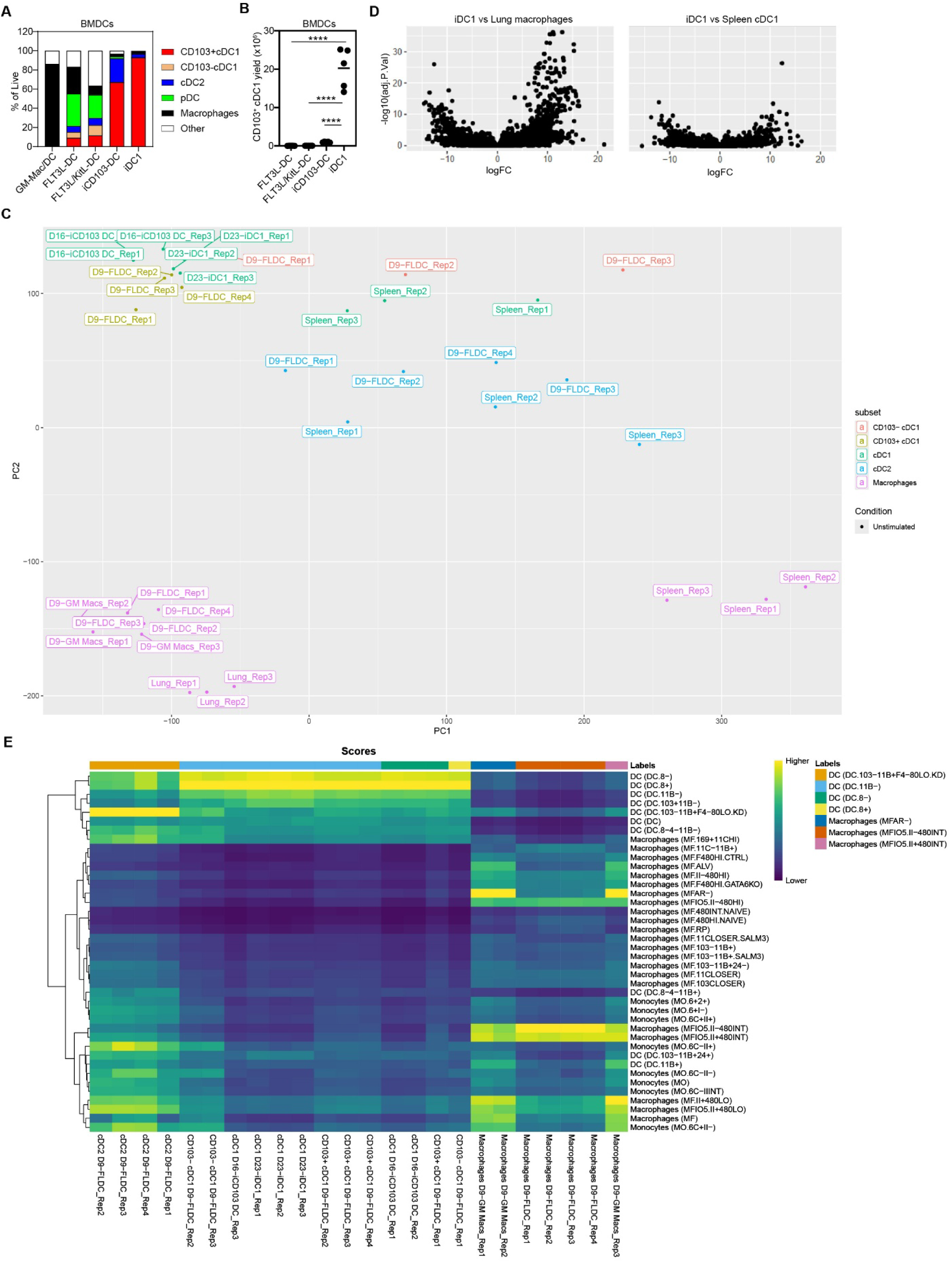
iDC1 cultures generate homogeneous CD103+ cDC1 with phenotypic and transcriptional features of the cDC1 lineage. (A, B) iDC1 were compared with other bone marrow-derived DC (BMDC) culture systems by flow cytometry. (A) Cellular composition. (B) Total numbers of CD103+ cDC1 generated per 1 x 106 bone marrow input cells. (C, D) RNA-seq analysis of DC and macrophage populations purified from indicated BMDC cultures and tissues. (C) Principal component analysis. (D) Volcano plots showing differentially expressed genes in iDC1 compared with lung macrophages (left) and splenic cDC1 (right). (E) Heatmap showing transcriptional similarity between BMDC subsets and reference populations from the ImmGen database. **** p<0.0001 (unpaired two-tailed Student’s t-test).

FLT3L-DC cultures were highly heterogeneous, comprising 15.2% cDC1, 6.2% cDC2, and 34.0% pDC (Figure 2A). cDC1 could be further subdivided into CD103+ and CD103- subsets (9.1% and 6.1%, respectively), both expressing CD24 and Xcr1 (Figure 2A and Supplementary Figure S2A). FLT3L-DC also contained 27.8% CD172a+CD64+ macrophage-like cells expressing F4/80 and CD11b with low I-A/I-E, as well as 16.8% cells that did not fall within cDC1, cDC2, or macrophage gates (Figure 2A and Supplementary Figure S2A). FLT3L/KitL cultures showed a broadly similar composition but with increased cDC1 frequencies and reduced pDC and macrophage populations (Figure 2A and Supplementary Figure S2A). Together, these findings demonstrate that FLT3L-based cultures generate a broad spectrum of DC and macrophage populations and therefore provide a heterogeneous system suited to study DC subset development and lineage relationships, but not selective cDC1 generation.

iCD103-DC cultures contained 67.2% CD103+ cDC1 and 24.7% cDC2, with minimal pDC (2.1%), macrophages (2.7%), and non-DC populations (3.3%), confirming enrichment for CD103+ cDC1 relative to FLT3L-based systems^16^ but retaining a residual cDC2 fraction. In contrast, iDC1 cultures contained 92.8% CD103+ cDC1, did not generate appreciable CD103-cDC1 populations, and included <3.5% cDC2, pDC, macrophages, or non-DCs, respectively (Figure 2A and Supplementary Figure S2A), demonstrating a marked increase in homogeneity relative to all other BMDC systems tested. CD103+ cDC1 from iDC1 cultures lacked F4/80 expression, expressed low CD11b, and exhibited higher I-A/I-E levels compared with CD172a+CD64+ cells from GM-Mac/DC and FLT3L-DC cultures (Supplementary Figure S2B), consistent with a *bona fide* cDC1 phenotype. In addition, iDC1 cultures produced substantially higher total numbers of CD103+ cDC1 than all other BMDC methods tested when calculated per 1 x 10^6^ input bone marrow cells (Figure 2B). Together, these findings demonstrate that iDC1 cultures markedly improve both cDC1 purity and scalability relative to existing BMDC systems. Thus, whereas FLT3L-based cultures provide heterogeneous systems for studying multiple DC subsets, iCD103-DC and iDC1 cultures progressively increase selective enrichment of *bona fide* cDC1, with iDC1 achieving substantially greater purity and scalability.

To further define the transcriptional relationship between iDC1, other BMDCs, tissue DCs, and macrophages, we sorted CD103+ cDC1 (from iDC1, iCD103-DC, and FLT3L-DC cultures), CD103- cDC1 (from FLT3L-DC cultures), cDC2 (from FLT3L-DC cultures and spleen), CD64+ cells (from GM-Mac/DC and FLT3L-DC cultures), and tissue cDC1 and macrophages (from spleen and lung) by flow cytometry, followed by RNA-seq. Principal component analysis showed that CD64+ cells from GM-Mac/DC and FLT3L-DC cultures clustered closely with lung alveolar macrophages and were clearly separated from iDC1 and cDC populations (Figure 2C, D). In contrast, iDC1 clustered with cDC1 populations, showing highest similarity to CD103+ cDC1 from iCD103-DC and FLT3L-DC cultures and close similarity to CD103- FLT3L-DC and splenic cDC1 (Figure 2C, D). cDC2 populations formed a distinct cluster separate from cDC1 while remaining within a broader cDC cluster, whereas splenic red pulp macrophages clustered away from cDC populations and were also distinct from lung alveolar macrophages.

We next compared the BMDC populations from our dataset with the ImmGen reference atlas using SingleR-based analysis to independently assess their transcriptional identity. The ImmGen reference contains 830 microarray samples representing 253 finely resolved immune cell subtypes. cDC1 populations from iDC1, iCD103-DC, and FLT3L-DC cultures showed highest similarity to the SingleR reference categories DC.8+ and DC.8-, with additional similarity to DC.11B- and DC.103+11B- (Figure 2E). In contrast, FLT3L-derived cDC2 showed highest similarity to DC.103-11B+F4-80LO.KD together with additional similarity to other CD11b-associated dendritic cell, monocyte, and macrophage reference categories. GM-CSF-derived and FLT3L-derived macrophages instead showed highest similarity to macrophage-associated reference categories and remained clearly separated from the cDC-associated reference branch. The preferential alignment of cDC1 populations with a restricted group of related reference categories, together with the clear separation of cDC2 and macrophage populations, further supports the conclusion that iDC1, iCD103-DC, and FLT3L-derived cDC1 share a common cDC1-associated transcriptional program despite their distinct culture conditions. iDC1 clustered within the broader cDC1 group together with iCD103-DC and FLT3L-derived cDC1 populations while remaining clearly distinct from cDC2 and macrophage lineages. These findings provide an independent reference-based validation that iDC1 closely resemble bona fide cDC1 at the transcriptional level.

Together, these data demonstrate that iDC1 cultures generate a highly homogeneous population of CD103+ cDC1 that closely aligns with the cDC1 lineage at both phenotypic and transcriptional levels while remaining clearly distinct from macrophage populations.

### iDC1 respond to innate stimulation and cross-present cell-associated antigen

To evaluate whether iDC1 respond to innate stimuli, particularly via TLR3, a cDC1 signature gene^39^, we stimulated iDC1 with TLR ligands and analyzed expression of co-stimulatory molecules by flow cytometry. iDC1 responded to LPS, poly(I:C), and CpG ODN1826 (CpG) stimulation by upregulating CD80, CD86, and CD40, whereas co-stimulation with poly(I:C) and CpG further increased CD80 and CD40 expression (Figure 3A, B). In contrast, CD103 expression remained largely unchanged across conditions (Figure 3A), suggesting that innate activation does not substantially alter the differentiated cDC1 phenotype of iDC1.

**Figure 3.**
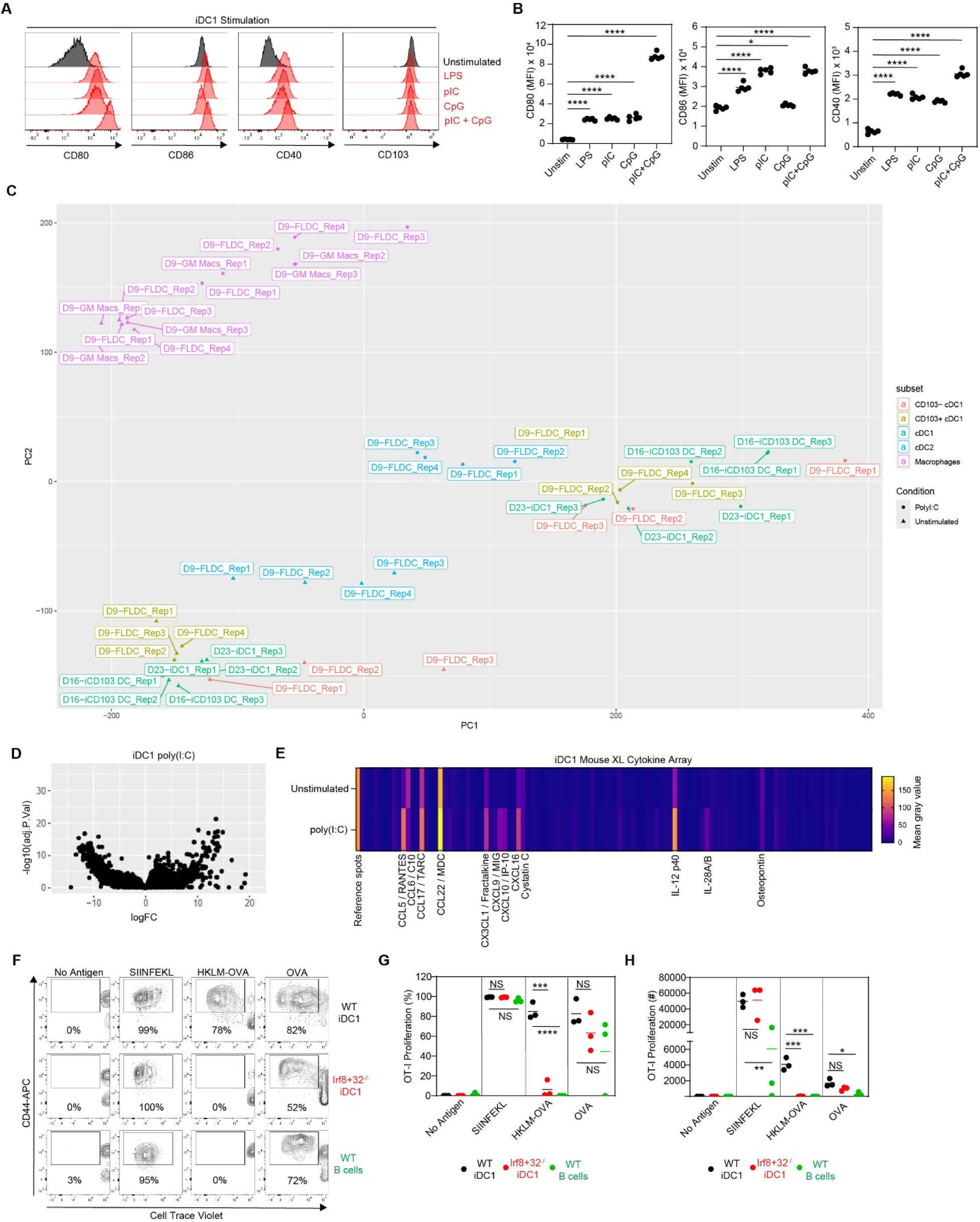
iDC1 respond to innate stimulation and exhibit functional properties of cDC1. (A, B) iDC1 were left unstimulated or stimulated overnight with LPS, poly(I:C) (pIC), CpG ODN1826 (CpG), or combined pIC and CpG. One of two independent experiments, each including 5 biological replicates, is shown. (A) Representative half-offset histograms showing CD80, CD86, CD40, and CD103 expression under unstimulated (black) and indicated stimulated conditions (red). (B) Quantification of mean fluorescence intensities (MFI). (C, D) RNA-seq analysis of DC and macrophage populations purified from indicated BMDC cultures that were either unstimulated or stimulated overnight with poly(I:C). (C) Principal component analysis. (D) Volcano plot showing differentially expressed genes. (E) Supernatants from unstimulated or poly(I:C)-stimulated iDC1 cultures were analyzed using a Mouse XL cytokine array. Heatmap showing mean gray values (MGV) for 111 targets under unstimulated and poly(I:C)-stimulated conditions. Targets detected above threshold (MGV > 20) in either condition are annotated. Data represent the average of duplicate spots per array from two independent experiments. (F-H) Cell Trace Violet (CTV)-labelled CD8+ OT-I T cells were co-cultured with equal numbers of WT iDC1, Irf8+32-/- iDC1, or B cells and were left unstimulated or stimulated with SIINFEKL peptide, cell-associated OVA (heat-killed Listeria monocytogenes expressing OVA; HKLM-OVA), or soluble OVA for 4 days. (F) Representative contour plots showing CTV dilution and CD44 expression of CD8+CD45.1+Vα2+ OT-I cells under the indicated conditions. (G, H) Quantification of proliferated (CTV-CD44+) OT-I cells as (G) percentages and (H) total cell numbers. Data are combined from three independent experiments. (B, G, H) * p<0.05, ** p<0.01, *** p<0.001, **** p<0.0001; NS, not statistically significant (two-tailed unpaired Student’s t-test).

To define the transcriptional response of iDC1 to poly(I:C) stimulation and compare it with other BMDC populations, we performed RNA-seq. Principal component analysis showed that macrophages clustered separately from cDC populations (Figure 3C). Following poly(I:C) stimulation, CD103+ cDC1 from iDC1, iCD103-DC, and FLT3L-DC cultures exhibited a more pronounced transcriptional shift compared with cDC2 and macrophages (Figure 3C). This enhanced transcriptional responsiveness is consistent with the specialized sensitivity of cDC1 to nucleic acid sensing pathways mediated by TLR3^2^. In iDC1, 3324 genes were differentially expressed following poly(I:C) stimulation (Figure 3D and Supplementary Data S1). Gene set enrichment analysis (GSEA) revealed that the top upregulated pathways were associated with interferon signaling, antiviral responses, inflammatory cytokine signaling, and innate immune activation, whereas downregulated pathways were enriched for cell-cycle, DNA replication, and proliferation-associated programs (Supplementary Data S2). Poly(I:C) stimulation therefore appears to shift iDC1 from a proliferative steady-state program toward an inflammatory effector state characterized by enhanced interferon signaling, innate activation, and cytokine-associated transcriptional responses.

To assess the functional consequences of poly(I:C) stimulation at the protein level, we profiled supernatants from unstimulated and poly(I:C)-stimulated iDC1 cultures using a Mouse XL cytokine array. iDC1 increased secretion of IL-12 p40 in response to stimulation (Figure 3E), a hallmark cytokine produced by activated cDC1^2–4^. Stimulated iDC1 also secreted chemokines including CCL5 (RANTES), CCL17 (TARC), CCL22 (MDC), CX3CL1 (Fractalkine), CXCL9 (MIG), CXCL10 (IP-10), and CXCL16, whereas secretion of CCL6 (C10) was reduced following poly(I:C) stimulation (Figure 3E). Several induced chemokines, including CXCL9, CXCL10, CCL5, and CX3CL1, are associated with recruitment and activation of cytotoxic lymphocytes and Th1-associated immune responses, consistent with established cDC1 functions^2–4^. Poly(I:C)-stimulated iDC1 also secreted IL-28A/B, whereas Cystatin C and Osteopontin were detected under both unstimulated and stimulated conditions (Figure 3E), further supporting acquisition of an inflammatory antiviral effector state.

We next compared WT iDC1, Irf8+32^-/-^ iDC1 (comprising cDC2 generated under identical conditions; Figure 1F), and B cells for their capacity to cross-present cell-associated and soluble OVA to OT-I T cells. No OT-I proliferation was observed in the absence of antigen, while all antigen-presenting cell (APC) types induced robust OT-I proliferation in response to SIINFEKL peptide (Figure 3F-H). Strikingly, WT iDC1, but not Irf8+32^-/-^ iDC1 or B cells, efficiently cross-presented cell-associated OVA (heat-killed *Listeria monocytogenes* expressing OVA; HKLM-OVA). In contrast, cross-presentation of soluble OVA was observed for both WT and Irf8+32^-/-^iDC1, although responses were significantly greater than those induced by B cells (Figure 3F-H). Together, these data demonstrate that iDC1 respond robustly to innate stimulation and exhibit key functional properties of *bona fide* cDC1, including inflammatory cytokine production and efficient cross-presentation of cell-associated antigen.

### GM-CSF and KitL regulate distinct stages of cDC1 generation in iDC1 cultures

To define how KitL and GM-CSF contribute to cDC1 generation during the 23-day iDC1 culture period, we compared five conditions: (1) iDC1 control, (2) no GM-CSF from day 9 to day 23, (3) no GM-CSF throughout the culture period, (4) no GM-CSF from day 0 to day 9, and (5) no KitL (extended iCD103-DC control; Figure 4A). iDC1 control cultures generated the highest purity of CD103+ cDC1, whereas purity was most reduced when GM-CSF was absent throughout the culture (condition 3) and was also significantly decreased when GM-CSF was removed from day 9 to day 23 (condition 2; Figure 4B). iDC1 control cultures also produced the highest total numbers of CD103+ cDC1, whereas all cytokine perturbation conditions markedly reduced cDC1 yields (Figure 4C). Notably, removal of GM-CSF during the early culture phase (condition 4) or omission of KitL (condition 5) substantially reduced total cDC1 numbers without markedly affecting cDC1 purity, suggesting that early KitL and GM-CSF primarily support efficient expansion of cDC1 precursors rather than later maintenance of cDC1 identity. Together, these results indicate that KitL and GM-CSF cooperate during the early phase of culture, whereas GM-CSF remains necessary during the later phase to maintain optimal cDC1 yield and purity, suggesting that GM-CSF directly regulates the maintenance and functional state of differentiated cDC1.

**Figure 4.**
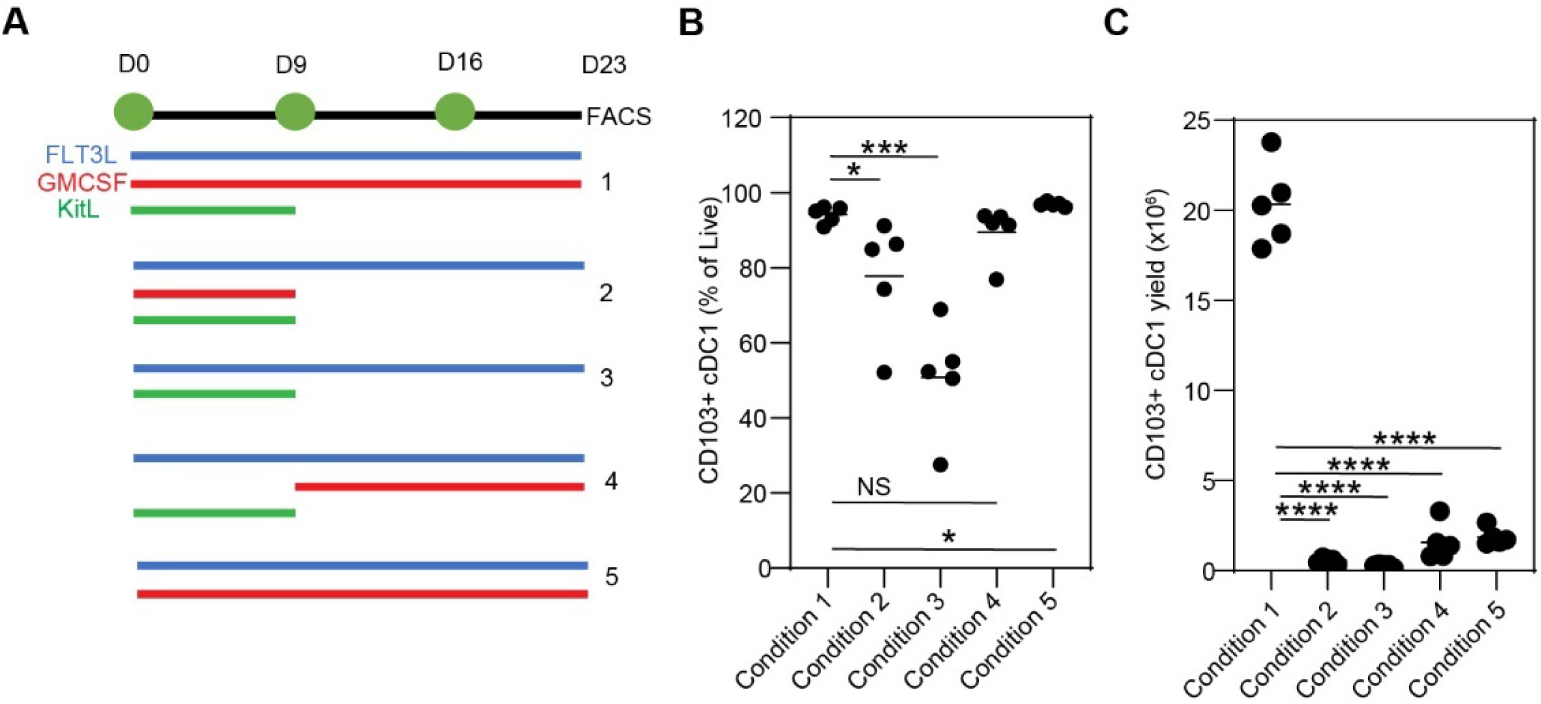
GM-CSF and KitL regulate distinct stages of cDC1 generation in iDC1 cultures. iDC1 and indicated variants were generated and analyzed by flow cytometry. (A) Overview of conditions tested. (B) Frequency of CD103+ cDC1 among live cells. (C) Total numbers of CD103+ cDC1 generated per 1 x 106 input bone marrow cells. One representative of two independent experiments, each including five biological replicates, is shown. * p<0.05, *** p<0.001, **** p<0.0001; NS, not statistically significant (two-tailed unpaired Student’s t-test).

### Proteomic analysis identifies GM-CSF–regulated pathways in iDC1

We next sought to define the GM-CSF-regulated pathways and biological processes sustaining the late phase of efficient cDC1 generation in iDC1 cultures. Because GM-CSF withdrawal could reduce cDC1 viability and therefore compromise proteomic resolution, we first tested whether Bcl-2 expression preserves cDC1 survival under these conditions. iDC1 cultures were therefore generated from Xcr1^Cre^ controls and Xcr1^Cre^ Rosa26^LSL-Bcl2-IRES-GFP^ bone marrow, resulting in Bcl-2 and GFP reporter expression in cDC1, followed by ∼21 h culture with FLT3L alone or FLT3L plus GM-CSF (Supplementary Figure S3A). Bcl-2 expression, indicated by GFP reporter expression, did not affect CD103+ cDC1 generation on day 23 and only modestly affected survival on day 24 in the continued presence of GM-CSF (Supplementary Figure S3B-E). In contrast, Bcl-2 markedly enhanced cDC1 survival following GM-CSF withdrawal, restoring viability to levels comparable with controls maintained in GM-CSF (Supplementary Figure S3B-E).

To identify GM-CSF-regulated pathways independently of acute effects on cell viability, iDC1 cultures generated from Xcr1^Cre^ Rosa26^LSL-Bcl2-IRES-GFP^ bone marrow were cultured for 20 h with FLT3L alone or FLT3L plus GM-CSF prior to proteomic analysis. GM-CSF significantly up-regulated 45 proteins and significantly down-regulated 12 proteins with a Log2FC > 0.6 (Figure 5B). In addition, GM-CSF significantly increased 268 and decreased 45 phosphorylation events (Figure 5C). Ingenuity pathway analysis identified prominent regulation of RHO GTPase signaling, cytoskeletal organization, membrane trafficking, and endocytosis pathways in both datasets (Figure 5D, E and Supplementary Data S3). These pathways are notable because cytoskeletal remodeling and membrane trafficking are central to DC migration, antigen uptake, and cross-presentation, processes previously linked to GM-CSF signaling^40–43^. Apoptotic execution, oxidative stress-, and metabolism-associated pathways were also significantly represented in both the global and phospho-proteomic datasets, whereas phospho-proteomic analysis more prominently identified pathways associated with cell-cycle progression, chromosomal replication, and transcriptional regulation (Figure 5D, E and Supplementary Data S3). Additional phospho-proteomic pathways included RNA Polymerase II transcription, DNA methylation and transcriptional repression signaling, and pathways linked to vesicular trafficking and cytoskeletal remodeling, suggesting that GM-CSF maintains a broadly active transcriptional and biosynthetic state in differentiated iDC1.

**Figure 5.**
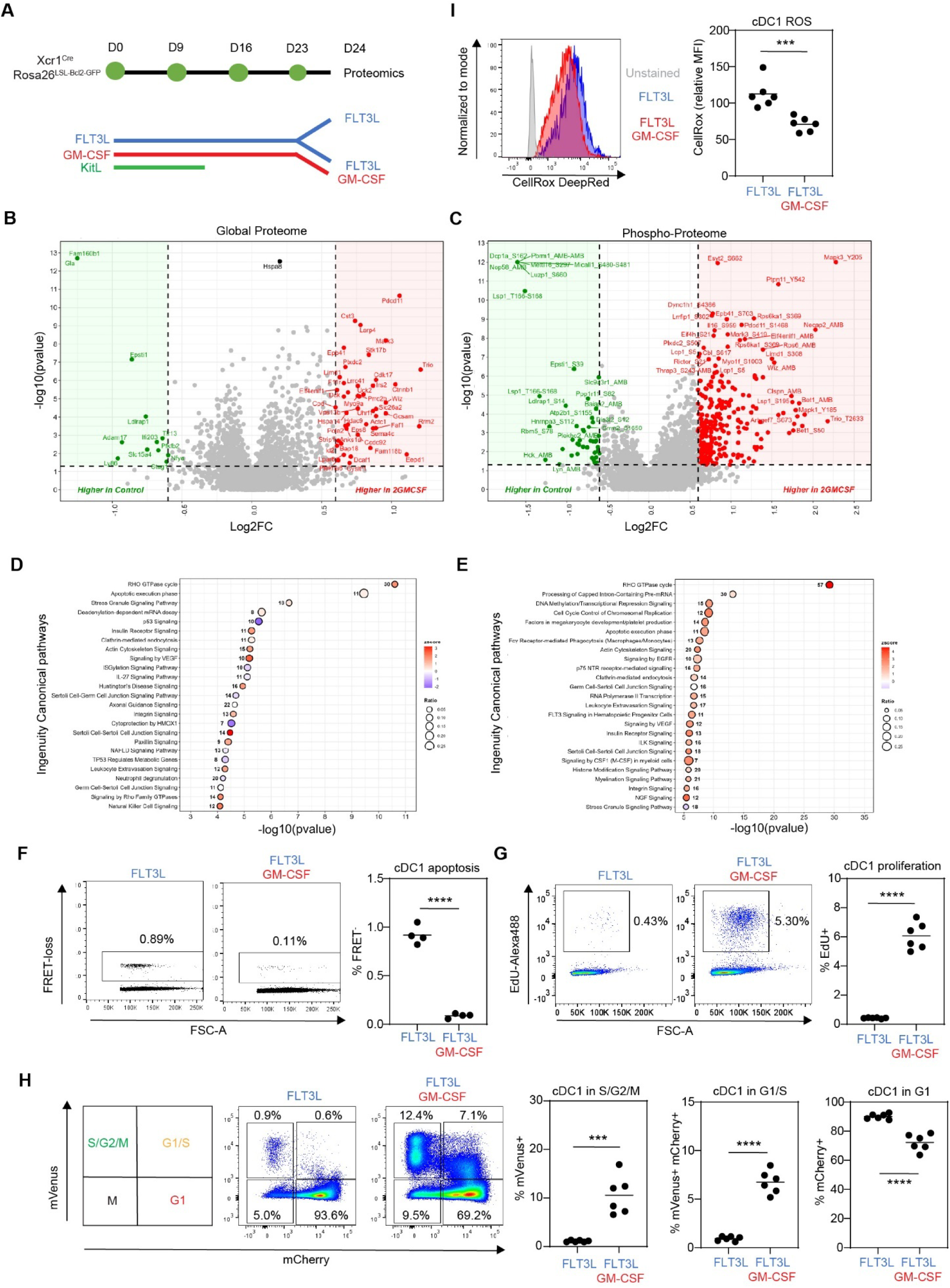
Proteomic and phospho-proteomic analyses identify GM-CSF-regulated signaling and biological programs in iDC1. iDC1 cultures were generated from the indicated genotypes for 23 days and subsequently cultured for an additional 20 - 21 h with FLT3L alone or FLT3L plus GM-CSF. (A–E) Proteomic and phospho-proteomic analyses. Xcr1Cre Rosa26LSL-Bcl2-IRES-GFP bone marrow was used to prevent apoptosis following GM-CSF withdrawal (Supplementary Figure S3). (A) Experimental scheme. (B, C) Volcano plots showing GM-CSF-regulated (B) proteins and (C) phosphorylation events. (D, E) Ingenuity pathway analysis of GM-CSF-regulated (D) proteins and (E) phosphorylation events. The top 25 pathways are shown. (F) Analysis of cDC1 apoptosis using Rosa26INDIA bone marrow. Representative dot plots on the left show the percentage of FRET- early apoptotic cells, with quantification shown on the right. (G) Analysis of EdU incorporation to identify cDC1 in S phase of the cell cycle. Representative contour plots on the left show FSC and EdU incorporation, with quantification of EdU+ cDC1 shown on the right. (H) Rosa26Fucci2aR bone marrow was used to assess cell-cycle stages. Representative contour plots on the left show percentages of cDC1 in G1, G1/S, and S/G2/M phases, with quantification shown on the right. (I) iDC1 were stained with CellRox FarRed dye to assess reactive oxygen species (ROS). Histograms on the left show CellRox staining of iDC1 cultured with FLT3L alone (blue), FLT3L plus GM-CSF (red), or unstained control (gray). CellRox mean fluorescence intensity (MFI) is quantified on the right. (F–I) Results are representative of, or combined from two independent experiments, each including 3–4 biological replicates. *** p<0.001, **** p<0.0001; NS, not significant (two-tailed unpaired Student’s t-test).

Notably, phosphorylation of Mapk3/Erk1 at the activating Y205 site represented one of the strongest GM-CSF-regulated phospho-events (Figure 5C), together with signaling changes consistent with engagement of MAPK-, PI3K/AKT-, mTOR-, and STAT5-associated pathways downstream of GM-CSF signaling^44^ (Supplementary Data S3). Additional regulated phosphosites included Ptpn11/SHP2 Y542 and Rps6ka1 S369, further supporting activation of canonical GM-CSF-associated MAPK signaling networks. Previous transcriptomic studies linked GM-CSF-responsive transcriptional programs to cDC1 survival and proliferation programs^45^. Our proteomic and phospho-proteomic analyses extend these findings by identifying coordinated regulation of signaling, cytoskeletal, membrane trafficking, oxidative stress, and transcriptional programs associated with maintenance of differentiated cDC1. Together, these data suggest that GM-CSF supports a proliferative, transcriptionally active, and survival-permissive state associated with extensive cytoskeletal and membrane trafficking programs in differentiated iDC1.

### GM-CSF promotes survival and proliferation of differentiated cDC1

To functionally validate the proteomic findings, we generated iDC1 cultures from Rosa26^INDIA^ mice, which sensitively report active caspase-3 through loss of FRET^21^. GM-CSF significantly suppressed FRET loss in cDC1 (Figure 5F), whereas this effect was less apparent in cDC2 within the same cultures, although low cDC2 numbers limited this analysis (Supplementary Figure S4A). We next quantified cells in S phase of the cell cycle by pulsing iDC1 cultures with EdU for 2 h followed by flow cytometric detection of EdU incorporation into DNA. GM-CSF significantly increased the percentage of cDC1 in S phase, whereas the effect was less pronounced in cDC2 (Figure 5G and Supplementary Figure S4B). As an independent approach to assess cell-cycle progression, we generated iDC1 cultures from Rosa26^Fucci2aR^ mice^25^, which distinguish G1, G1/S, and S/G2/M phases through mCherry and mVenus expression. GM-CSF markedly increased the percentages of cDC1 in G1/S and S/G2/M phases while decreasing the proportion of cells in G1 (Figure 5H). These changes were less pronounced or absent in cDC2 (Supplementary Figure S4C). We also evaluated production of reactive oxygen species (ROS) in live cDC1 by staining with the cell-permeable dye CellRox FarRed. GM-CSF significantly reduced ROS production (Figure 5I), consistent with a role for GM-CSF in maintaining redox homeostasis and limiting apoptosis. Together, these experiments validate the major biological programs identified by the proteomic analyses and demonstrate that GM-CSF suppresses apoptosis, limits oxidative stress, and promotes cDC1 cell-cycle progression, thereby contributing mechanistically to the preferential outgrowth and sustained maintenance of cDC1 during the later stages of iDC1 culture.

### STAT5- and BRD4-associated regulatory programs contribute to efficient iDC1 generation

Transcription factor target enrichment analysis implicated STAT5-, IRF8/BATF-, MYC-, and BRD4-associated regulatory programs among GM-CSF-responsive proteins (Supplementary Data S4). STAT5 is a canonical downstream mediator of GM-CSF signaling^43^, whereas MYC-associated programs have previously been linked to GM-CSF-dependent cDC1 proliferation and survival^45^. BRD4 regulates transcriptional elongation and MYC-associated transcriptional programs linked to highly active cellular states^23,46^. We therefore investigated whether STAT5 and BRD4 contribute to efficient cDC1 generation in iDC1 cultures.

To assess the role of STAT5, iDC1 cultures were generated from Xcr1^Cre^ Stat5a/b^fl/fl^ bone marrow and corresponding controls. Functional loss of STAT5 signaling in Xcr1^Cre^Stat5a/b^fl/fl^ cDC1 was confirmed by flow cytometric analysis of phosphorylated STAT5 (p-STAT5) following GM-CSF stimulation, which showed markedly reduced p-STAT5 induction relative to stimulated controls (Supplementary Figure S5A). Although STAT5-deficient iDC1 cultures retained comparably high cDC1 frequencies within the gated cDC compartment (Figure 6A), overall cDC1 purity was significantly reduced relative to control iDC1 cultures (Figure 6B), accompanied by modest increases in cDC2 and macrophage populations (Figure 6C). More importantly, total CD103+ cDC1 yields were dramatically reduced in Xcr1^Cre^Stat5a/b^fl/fl^ iDC1 cultures compared with WT controls (Figure 6D), indicating that STAT5 contributes to efficient cDC1 generation and maintenance in iDC1 cultures. Residual STAT5-deficient cDC1 also exhibited significantly reduced CD11c and CD103 expression (Figure 6A and Supplementary Figure S5B-D), consistent with previous studies linking GM-CSF signaling to regulation of the CD103+ cDC1 phenotype^40^. To determine whether STAT5 contributes to GM-CSF-mediated regulation of cDC1 proliferation and survival, day 23 iDC1 were cultured with FLT3L plus GM-CSF for 21 h prior to analysis of EdU incorporation and active caspase-3 (aCasp3). Xcr1^Cre^ Stat5a/b^fl/fl^ cDC1 exhibited significantly reduced proportions of EdU+ cells in S phase of the cell cycle (Figure 6E, F), accompanied by significantly increased frequencies of aCasp3+ apoptotic cells (Figure 6G, H). Together, these findings support a role for STAT5 downstream of GM-CSF signaling in promoting cDC1 proliferation and survival, extending previous observations that conditional STAT5 deletion reduces CD103+ DC populations in non-lymphoid tissues^47,48^.

**Figure 6.**
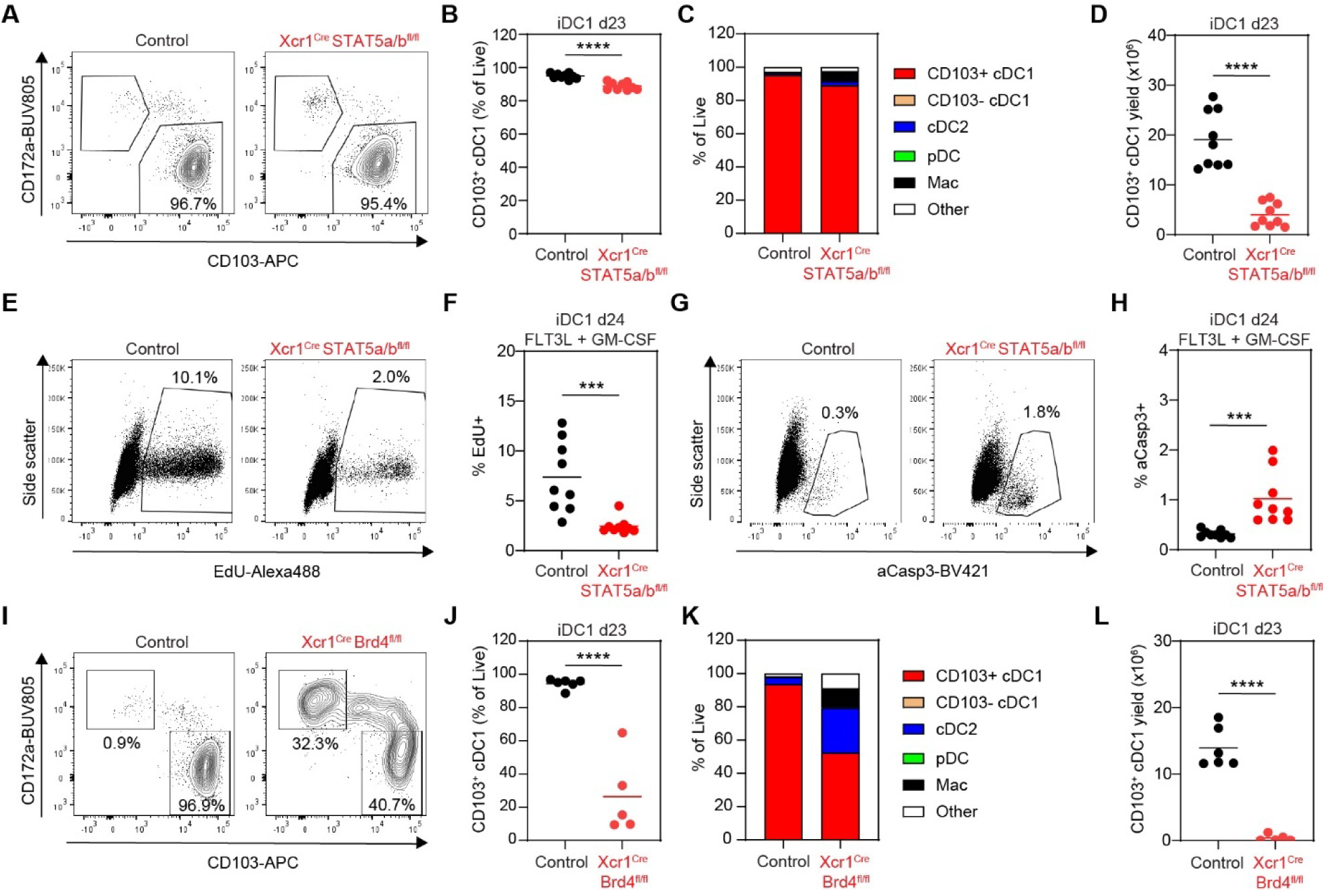
STAT5- and BRD4-associated regulatory programs contribute to efficient iDC1 generation. iDC1 cultures were generated from the indicated genotypes for 23 days and analyzed by flow cytometry on day 23 or cultured with FLT3L plus GM-CSF for 20–21 h, pulsed with EdU during the final 2 h of culture, and analyzed by flow cytometry on day 24. (A-H) Analysis of (A-D) day 23 or (E-H) day 24 STAT5-deficient iDC1 cultures. Results are combined from two independent experiments, each including 4–5 biological replicates per genotype. (A) Representative contour plots showing CD103 and CD172a expression and percentages of cDC1 among CD11c+CD45R/B220-CD64- cDC. (B) Quantification of CD103+ cDC1 purity. (C) Average composition of iDC1 cultures across replicates and experiments. (D) Total numbers of CD103+ cDC1 generated per 1 x 106 input bone marrow cells. (E) Representative dot plots showing EdU staining, side scatter, and percentages of EdU+ cDC1. (F) Quantification of EdU+ cDC1. (G) Representative dot plots showing active caspase-3 (aCasp3) staining, side scatter, and percentages of aCasp3+ cDC1. (H) Quantification of aCasp3+ cDC1. (I-L) Analysis of day 23 BRD4-deficient iDC1 cultures. Results are combined from three independent experiments, each including 1–4 biological replicates per genotype. (I) Representative contour plots showing CD103 and CD172a expression and percentages of cDC1 and cDC2 among CD11c+CD45R/B220-CD64- cDC. (J) Quantification of CD103+ cDC1 purity. (K) Average composition of iDC1 cultures across replicates and experiments. (L) Total numbers of CD103+ cDC1 generated per 1 x 106 input bone marrow cells. *** p<0.001, **** p<0.0001 (two-tailed unpaired Student’s t-test).

Although the BRD4 program was identified (Supplementary Data S4), its role in cDC1 biology remains poorly defined. We next assessed the role of BRD4 by generating iDC1 cultures from Xcr1^Cre^Brd4^fl/fl^ bone marrow. A fraction of control cDC1 expressed BRD4 which was efficiently ablated in Xcr1^Cre^Brd4^fl/fl^ cDC1, while Brd4 expression in cDC2 was not affected (Supplementary Figure S5E). Brd4 deficiency caused pronounced CD11c downregulation and significant CD103 upregulation (Figure 6I and Supplementary Figure S5F-H). These findings suggest that BRD4 regulates multiple aspects of the differentiated cDC1 phenotype. Additionally, CD103+ cDC1 purity was significantly reduced (Figure 6I, J), mainly due to increases in cDC2, macrophages, and cells not falling within any of the gates (Figure 6K). Most importantly, BRD4 deficiency almost completely abrogated cDC1 generation in iDC1 cultures (Figure 6L). These findings identify STAT5- and BRD4-associated regulatory programs as important regulators of efficient iDC1 generation.

## Conclusion

We developed a scalable recombinant cytokine-based BMDC system termed iDC1 that enables efficient generation of highly pure +32 kb *Irf8* enhancer-dependent CD103+ cDC1. Compared with existing BMDC methods, iDC1 cultures markedly improve cDC1 purity and yield while minimizing contamination by cDC2, macrophages and other non-DC populations. iDC1 also exhibit phenotypic, transcriptional, and functional properties characteristic of bona fide cDC1, including robust innate responsiveness, IL-12 production, and efficient cross-presentation of cell-associated antigen.

Mechanistically, our findings identify distinct temporal requirements for KitL and GM-CSF during cDC1 generation and demonstrate that GM-CSF maintains differentiated cDC1 through coordinated regulation of survival, proliferation, oxidative stress, signaling, cytoskeletal remodeling, membrane trafficking, and transcriptional programs. Proteomic and phospho-proteomic analyses further identified STAT5- and BRD4-associated regulatory programs as important contributors to efficient iDC1 generation.

The iDC1 platform should facilitate future studies of cDC1 development, signaling, metabolism, antigen processing, and immunotherapy. In particular, the ability to generate large numbers of highly pure cDC1 using recombinant cytokines may support pre-clinical studies investigating cDC1-based vaccination, tumor immunity, and engineered dendritic-cell therapies.

## Acknowledgements

We thank Dr. Dinah Singer for critical reading of the manuscript; Jeffrey Chiang and Jie Mu for technical support; Assiatu Crossman, Kheem Bisht, Don Plugge, William Hajjar, and Tengfei Zhang for flow cytometry support; all staff at the NCI Frederick and NCI Bethesda animal facilities for their critical help. During the preparation of this work, the authors used Claude and ChatGPT provided by the U.S. Department of Health and Human Services in order to optimize language. After using these tools, the authors reviewed and edited the content as needed and take full responsibility for the content of the publication. The content of this publication does not necessarily reflect the views or policies of the Department of Health and Human Services, nor does mention of trade names, commercial products, or organizations imply endorsement by the U.S. Government.

## Author’s Contribution

Sharma S, Flynn F, Capaldo B, Holewinski R, Chen Q, Meerzaman D, Andresson T, Mayer CT:

Data acquisition, analysis, or interpretation

Sharma S, Mayer CT: Manuscript revision

Mayer CT: Article conception and design

## Ethical Approval

All procedures were approved by the NCI Animal Care and Use Committee (ACUC) and conformed with federal regulatory requirements and standards. The intramural NIH ACU program is accredited by AAALAC International.

## Consent to Participate

Not applicable

## Consent for Publication

Not applicable

## Availability of Data and Materials

RNA-seq data are available at NCBI Accession PRJNA1474846. Proteomics data are available at MassIVE Accession MSV000101983.

## References

1 Durai, V. & Murphy, K. M. Functions of Murine Dendritic Cells. Immunity 45, 719–736 (2016). 10.1016/j.immuni.2016.10.010

2 Hildner, K. et al. Batf3 deficiency reveals a critical role for CD8alpha+ dendritic cells in cytotoxic T cell immunity. Science 322, 1097–1100 (2008). 10.1126/science.1164206

3 Mashayekhi, M. et al. CD8alpha(+) dendritic cells are the critical source of interleukin-12 that controls acute infection by Toxoplasma gondii tachyzoites. Immunity 35, 249–259 (2011). 10.1016/j.immuni.2011.08.008

4 Koblansky, A. A. et al. Recognition of profilin by Toll-like receptor 12 is critical for host resistance to Toxoplasma gondii. Immunity 38, 119–130 (2013). 10.1016/j.immuni.2012.09.016

5 Canton, J. et al. The receptor DNGR-1 signals for phagosomal rupture to promote cross-presentation of dead-cell-associated antigens. Nat Immunol 22, 140–153 (2021). 10.1038/s41590-020-00824-x

6 Tussiwand, R. et al. Compensatory dendritic cell development mediated by BATF-IRF interactions. Nature 490, 502–507 (2012). 10.1038/nature11531

7 Glasmacher, E. et al. A genomic regulatory element that directs assembly and function of immune-specific AP-1-IRF complexes. Science 338, 975–980 (2012). 10.1126/science.1228309

8 Durai, V. et al. Cryptic activation of an Irf8 enhancer governs cDC1 fate specification. Nat Immunol 20, 1161–1173 (2019). 10.1038/s41590-019-0450-x

9 Helft, J. et al. GM-CSF Mouse Bone Marrow Cultures Comprise a Heterogeneous Population of CD11c(+)MHCII(+) Macrophages and Dendritic Cells. Immunity 42, 1197–1211 (2015). 10.1016/j.immuni.2015.05.018

10 Inaba, K. et al. Generation of large numbers of dendritic cells from mouse bone marrow cultures supplemented with granulocyte/macrophage colony-stimulating factor. J Exp Med 176, 1693–1702 (1992). 10.1084/jem.176.6.1693

11 Lutz, M. B. et al. An advanced culture method for generating large quantities of highly pure dendritic cells from mouse bone marrow. J Immunol Methods 223, 77–92 (1999). 10.1016/s0022-1759(98)00204-x

12 Brasel, K., De Smedt, T., Smith, J. L. & Maliszewski, C. R. Generation of murine dendritic cells from flt3-ligand-supplemented bone marrow cultures. Blood 96, 3029–3039 (2000).

13 Naik, S. H., O’Keeffe, M., Proietto, A., Shortman, H. H. & Wu, L. CD8+, CD8-, and plasmacytoid dendritic cell generation in vitro using flt3 ligand. Methods Mol Biol 595, 167–176 (2010). 10.1007/978-1-60761-421-0_10

14 Ashayeripanah, M. et al. Interleukin 4 selectively expands functional type 1 conventional dendritic cells from bone marrow progenitors. Cell Rep 45, 116772 (2026). 10.1016/j.celrep.2025.116772

15 Kirkling, M. E. et al. Notch Signaling Facilitates In Vitro Generation of Cross-Presenting Classical Dendritic Cells. Cell Rep 23, 3658–3672 e3656 (2018). 10.1016/j.celrep.2018.05.068

16 Mayer, C. T. et al. Selective and efficient generation of functional Batf3-dependent CD103+ dendritic cells from mouse bone marrow. Blood 124, 3081–3091 (2014). 10.1182/blood-2013-12-545772

17 Ou, F. et al. Enhanced in vitro type 1 conventional dendritic cell generation via the recruitment of hematopoietic stem cells and early progenitors by Kit ligand. Eur J Immunol 53, e2250201 (2023). 10.1002/eji.202250201

18 Ferris, S. T. et al. cDC1 prime and are licensed by CD4(+) T cells to induce anti-tumour immunity. Nature 584, 624–629 (2020). 10.1038/s41586-020-2611-3

19 Schwenk, F., Baron, U. & Rajewsky, K. A cre-transgenic mouse strain for the ubiquitous deletion of loxP-flanked gene segments including deletion in germ cells. Nucleic Acids Res 23, 5080–5081 (1995). 10.1093/nar/23.24.5080

20 Knittel, G. et al. B-cell-specific conditional expression of Myd88p.L252P leads to the development of diffuse large B-cell lymphoma in mice. Blood 127, 2732–2741 (2016). 10.1182/blood-2015-11-684183

21 Mayer, C. T. et al. The microanatomic segregation of selection by apoptosis in the germinal center. Science 358 (2017). 10.1126/science.aao2602

22 Cui, Y. et al. Inactivation of Stat5 in mouse mammary epithelium during pregnancy reveals distinct functions in cell proliferation, survival, and differentiation. Mol Cell Biol 24, 8037–8047 (2004). 10.1128/MCB.24.18.8037-8047.2004

23 Devaiah, B. N. et al. BRD4 is a histone acetyltransferase that evicts nucleosomes from chromatin. Nat Struct Mol Biol 23, 540–548 (2016). 10.1038/nsmb.3228

24 Hogquist, K. A. et al. T cell receptor antagonist peptides induce positive selection. Cell 76, 17–27 (1994). 10.1016/0092-8674(94)90169-4

25 Mort, R. L. et al. Fucci2a: a bicistronic cell cycle reporter that allows Cre mediated tissue specific expression in mice. Cell Cycle 13, 2681–2696 (2014). 10.4161/15384101.2015.945381

26 von Boehmer, L. et al. Sequencing and cloning of antigen-specific antibodies from mouse memory B cells. Nat Protoc 11, 1908–1923 (2016). 10.1038/nprot.2016.102

27 Simpson, M. J. et al. Peripheral apoptosis and limited clonal deletion during physiologic murine B lymphocyte development. Nat Commun 15, 4691 (2024). 10.1038/s41467-024-49062-x

28 Soneson, C., Love, M. I. & Robinson, M. D. Differential analyses for RNA-seq: transcript-level estimates improve gene-level inferences. F1000Res 4, 1521 (2015). 10.12688/f1000research.7563.2

29 Chen, Y., Chen, L., Lun, A. T. L., Baldoni, P. L. & Smyth, G. K. edgeR v4: powerful differential analysis of sequencing data with expanded functionality and improved support for small counts and larger datasets. Nucleic Acids Res 53 (2025). 10.1093/nar/gkaf018

30 Chen, Y., Lun, A. T. & Smyth, G. K. From reads to genes to pathways: differential expression analysis of RNA-Seq experiments using Rsubread and the edgeR quasi-likelihood pipeline. F1000Res 5, 1438 (2016). 10.12688/f1000research.8987.2

31 Ritchie, M. E. et al. limma powers differential expression analyses for RNA-sequencing and microarray studies. Nucleic Acids Res 43, e47 (2015). 10.1093/nar/gkv007

32 Hoffman, G. E. & Roussos, P. Dream: powerful differential expression analysis for repeated measures designs. Bioinformatics 37, 192–201 (2021). 10.1093/bioinformatics/btaa687

33 Hoffman, G. E. & Schadt, E. E. variancePartition: interpreting drivers of variation in complex gene expression studies. BMC Bioinformatics 17, 483 (2016). 10.1186/s12859-016-1323-z

34 Xu, S. et al. Using clusterProfiler to characterize multiomics data. Nat Protoc 19, 3292–3320 (2024). 10.1038/s41596-024-01020-z

35 Aran, D. et al. Reference-based analysis of lung single-cell sequencing reveals a transitional profibrotic macrophage. Nat Immunol 20, 163–172 (2019). 10.1038/s41590-018-0276-y

36 Kim, J. et al. Lysine methyltransferase Kmt2d regulates naive CD8(+) T cell activation-induced survival. Front Immunol 13, 1095140 (2022). 10.3389/fimmu.2022.1095140

37 Becattini, S. et al. Enhancing mucosal immunity by transient microbiota depletion. Nat Commun 11, 4475 (2020). 10.1038/s41467-020-18248-4

38 Sathaliyawala, T. et al. Mammalian target of rapamycin controls dendritic cell development downstream of Flt3 ligand signaling. Immunity 33, 597–606 (2010). 10.1016/j.immuni.2010.09.012

39 Miller, J. C. et al. Deciphering the transcriptional network of the dendritic cell lineage. Nat Immunol 13, 888–899 (2012). 10.1038/ni.2370

40 Zhan, Y. et al. GM-CSF increases cross-presentation and CD103 expression by mouse CD8(+) spleen dendritic cells. Eur J Immunol 41, 2585–2595 (2011). 10.1002/eji.201141540

41 Sathe, P. et al. The acquisition of antigen cross-presentation function by newly formed dendritic cells. J Immunol 186, 5184–5192 (2011). 10.4049/jimmunol.1002683

42 de Brito, C. et al. CpG promotes cross-presentation of dead cell-associated antigens by pre-CD8alpha+ dendritic cells [corrected]. J Immunol 186, 1503–1511 (2011). 10.4049/jimmunol.1001022

43 Shiomi, A. & Usui, T. Pivotal roles of GM-CSF in autoimmunity and inflammation. Mediators Inflamm 2015, 568543 (2015). 10.1155/2015/568543

44 van de Laar, L., Coffer, P. J. & Woltman, A. M. Regulation of dendritic cell development by GM-CSF: molecular control and implications for immune homeostasis and therapy. Blood 119, 3383–3393 (2012). 10.1182/blood-2011-11-370130

45 Kc, W. et al. L-Myc expression by dendritic cells is required for optimal T-cell priming. Nature 507, 243–247 (2014). 10.1038/nature12967

46 Devaiah, B. N. et al. MYC protein stability is negatively regulated by BRD4. Proc Natl Acad Sci U S A 117, 13457–13467 (2020). 10.1073/pnas.1919507117

47 Eddy, W. E. et al. Stat5 Is Required for CD103(+) Dendritic Cell and Alveolar Macrophage Development and Protection from Lung Injury. J Immunol 198, 4813–4822 (2017). 10.4049/jimmunol.1601777

48 Li, H. S. et al. The signal transducers STAT5 and STAT3 control expression of Id2 and E2-2 during dendritic cell development. Blood 120, 4363–4373 (2012). 10.1182/blood-2012-07-441311

