## Supplementary Information for "Scalable generation of pure CD103+ cDC1 from iDC1 cultures"

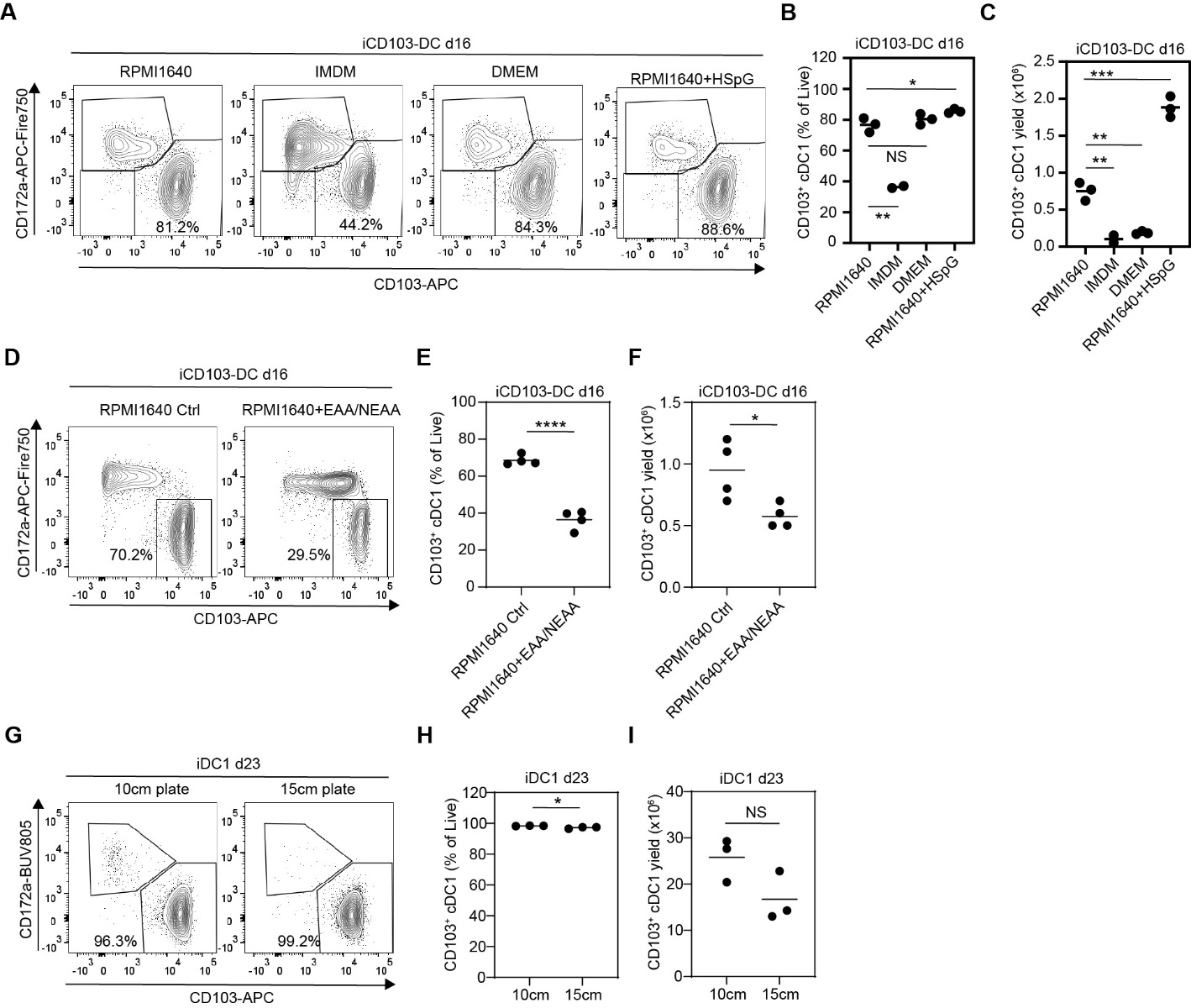

**Supplementary Figure S1. Effects of culture media composition and plate size on CD103^+^ cDC1 production.**

(A–F) iCD103-DC cultures were established in RPMI 1640 or IMDM or DMEM medium supplemented with 10% heat-inactivated FBS, 1X Penicillin/Streptomycin, and 1X 2-Mercaptoethanol (RPMI1640 Ctrl), or additionally supplemented with 1X MEM Amino Acids and 1X MEM Non-Essential Amino Acids (RPMI1640 + EAA/NEAA), or with 25 mM HEPES, 1 mM sodium pyruvate, and 2 mM Glutamax (RPMI1640 + HSpG). Cells were analyzed by flow cytometry on day 16. (A, D) Representative contour plots of CD103 versus CD172a expression among CD11c+B220- cDC. (B, E) Frequency of CD103+ cDC1 among live cells. (C, F) Total number of CD103+ cDC1 per 1 x 10^6^ input bone marrow cells. One of two representative experiments, each including 3–4 biological replicates, is shown. (G–I) iDC1 cultures were established in 10 cm or 15 cm plates with culture volumes scaled according to surface area, while maintaining constant cell and cytokine concentrations. (G) Representative contour plots of CD103 versus CD172a expression among CD11c+B220- cDC. (H) Frequency of CD103+ cDC1 among live cells. (I) Total number of CD103+ cDC1 per 1 x 10^6^ input bone marrow cells. One of three representative experiments, each including 1–4 biological replicates, is shown. * p<0.05, ** p<0.01, *** p<0.001, **** p<0.0001, NS = not statistically significant (unpaired two-tailed Student’s t-test).

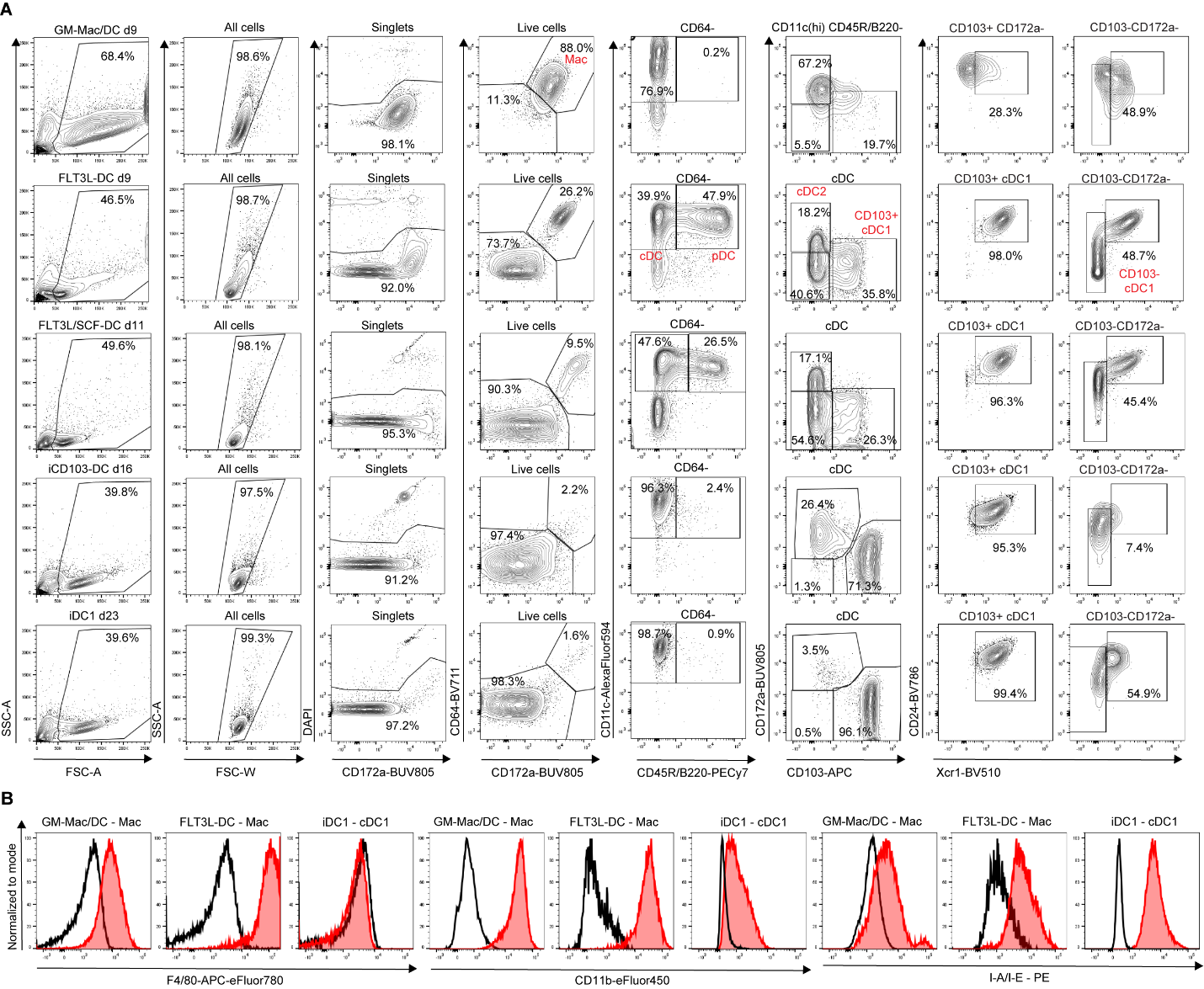

**Supplementary Figure S2. BMDC gating strategy and surface marker expression.** (A) Common flow cytometry gating strategy applied to the indicated BMDC culture systems. (B) Expression of F4/80, CD11b, and MHC-II among indicated subsets (red), overlaid with unstained controls (black).

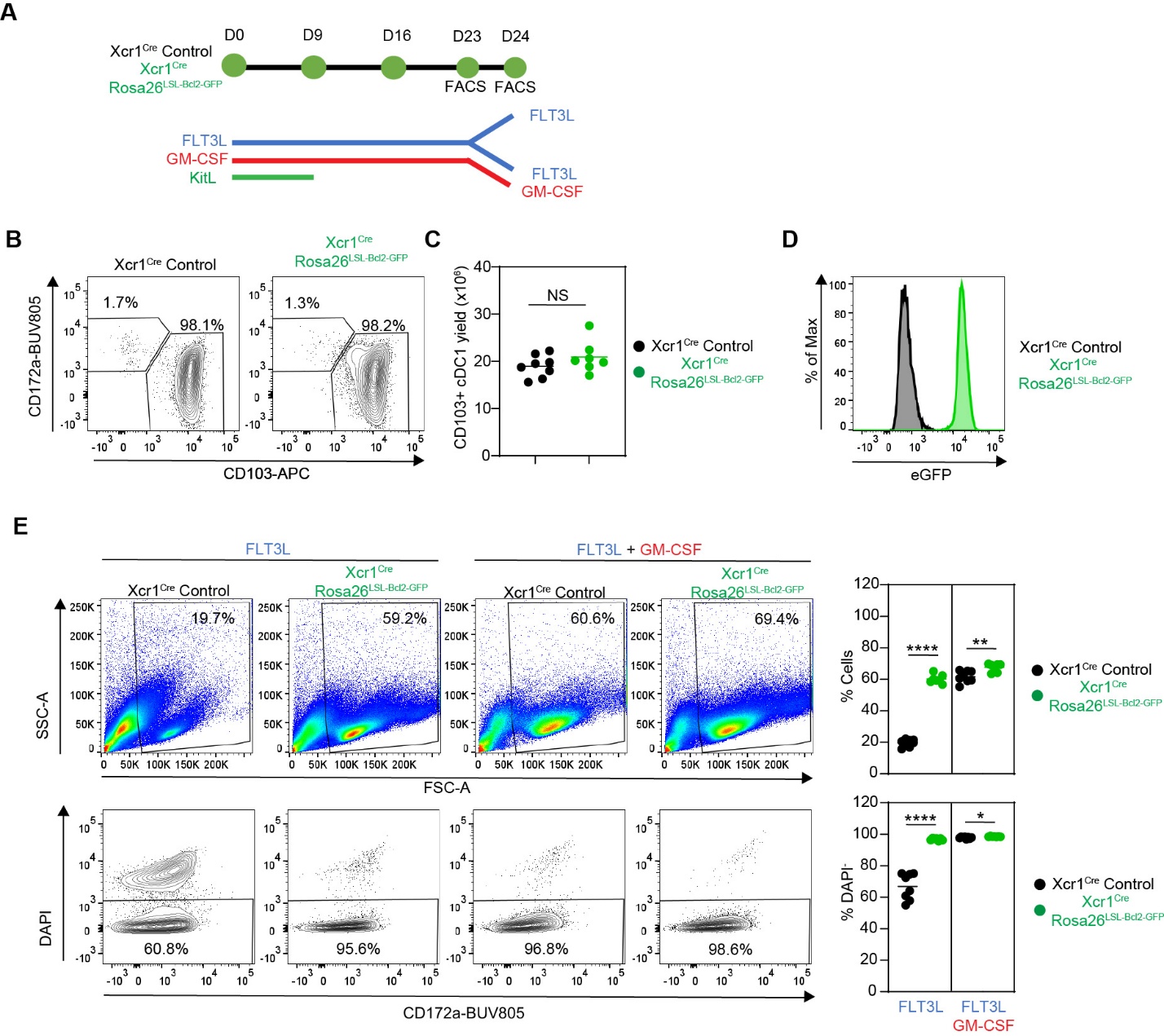

**Supplementary Figure S3. Bcl-2 expression does not affect iDC1 generation but promotes CD103^+^ cDC1 survival following GM-CSF withdrawal.** iDC1 cultures were generated from Xcr1^Cre^ control and Xcr1^Cre^ Rosa26^LSL-Bcl2-IRES-GFP^ bone marrow. On day 23, cells were either analyzed by flow cytometry or cultured for an additional 21 h with FLT3L alone or FLT3L plus GM-CSF prior to flow cytometric analysis on day 24. (A) Experimental scheme. (B-D) Day 23 analysis. (B) Representative contour plots showing CD103 and CD172a expression among CD11c^+^CD45R/B220^-^CD64^-^ cDC from the indicated genotypes. (C) Total numbers of CD103^+^ cDC1 generated per 1 x 10^6^ input bone marrow cells according to genotype. (D) Representative histogram shows GFP expression in CD103^+^ cDC1 from a Xcr1^Cre^ Rosa26^LSL-Bcl2-IRES-GFP^ mouse (green) relative to Xcr1^Cre^ control (black). (E) Day 24 analysis following GM-CSF withdrawal or continued GM-CSF exposure. Representative pseudocolor and contour plots showing forward and side scatter or CD172a expression and DAPI staining for identification of total and live cells under the indicated conditions. Graphs on the right show quantification. (D, E) Results are combined from two independent experiments, each including 3-4 biological replicates per genotype. * p<0.05, ** p<0.01, **** p<0.0001, NS = not statistically significant (unpaired two-tailed Student’s t-test).

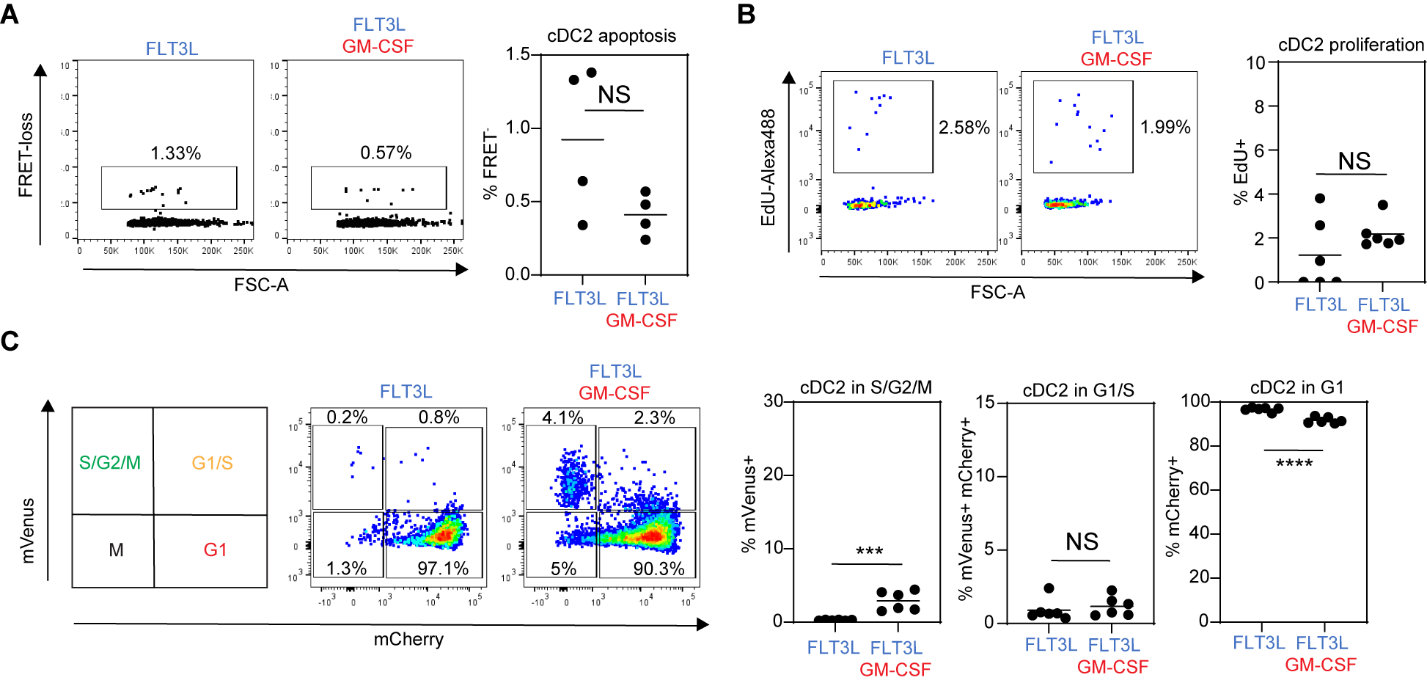

**Supplementary Figure S4. Analysis of cDC2 survival and proliferation in iDC1 cultures following GM-CSF withdrawal.** iDC1 cultures were generated from the indicated genotypes for 23 days and subsequently cultured for 20-21 h with FLT3L alone or FLT3L plus GM-CSF. (A) Analysis of cDC2 apoptosis using Rosa26^INDIA^ bone marrow. Representative dot plots on the left show the percentage of FRET- early apoptotic cells, with quantification shown on the right. (B) Analysis of EdU incorporation to identify cDC2 in S phase of the cell cycle. Representative contour plots on the left show FSC and EdU incorporation, with quantification of EdU+ cDC2 shown on the right. (C) Rosa26^Fucci2aR^ bone marrow was used to assess cell-cycle stages. Representative contour plots on the left show percentages of cDC2 in G1, G1/S, and S/G2/M phases, with quantification shown on the right. (A-C) Results are representative of, or combined from two independent experiments, each including 3–4 biological replicates. *** p<0.001, **** p<0.0001; NS, not significant (two-tailed unpaired Student’s *t*-test).

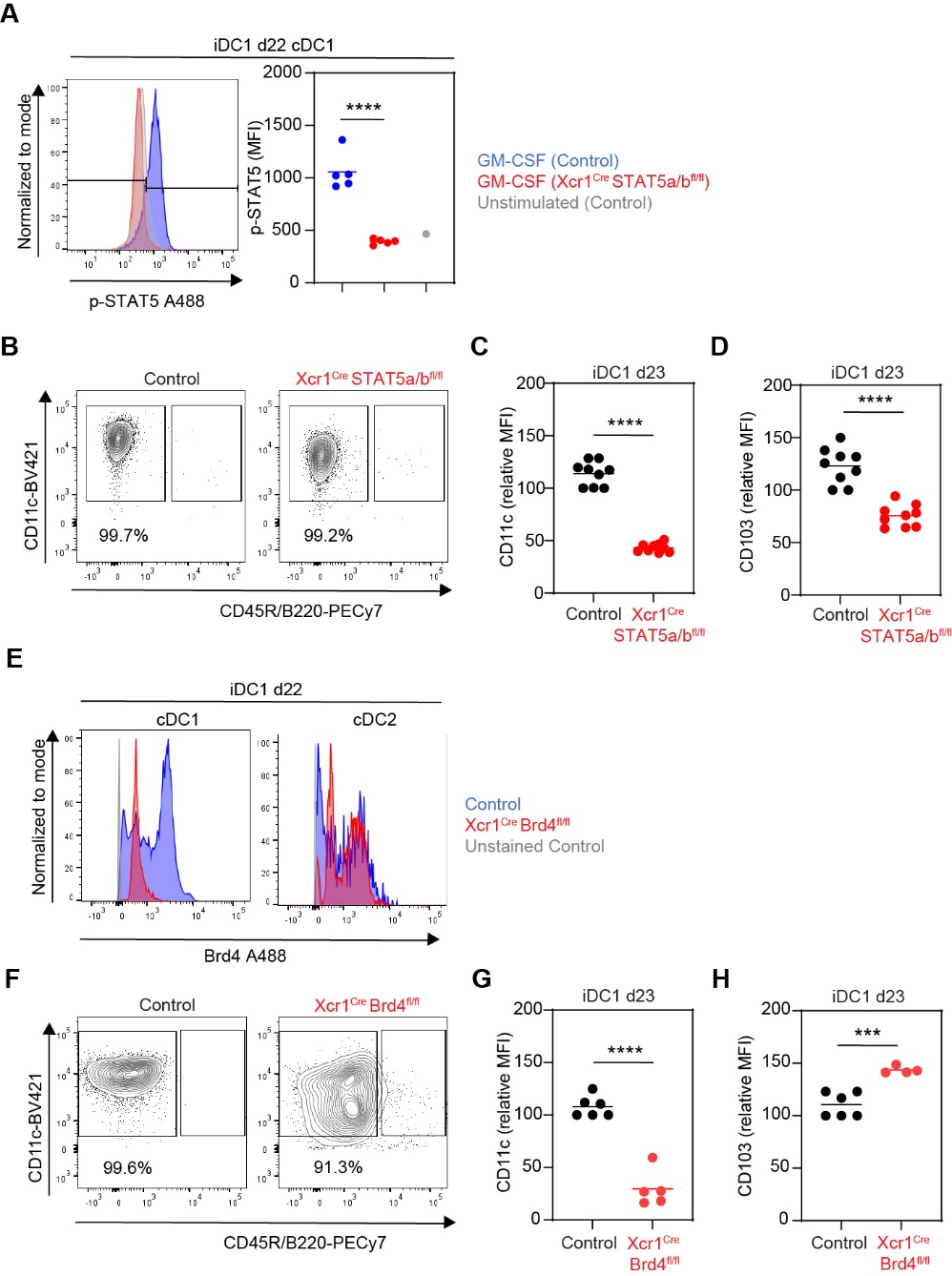

**Supplementary Figure S5. Validation of STAT5 and BRD4 deletion and effects on surface phenotype in iDC1 cultures.** iDC1 cultures were generated from the indicated genotypes and analyzed by flow cytometry on day 22 or day 23. (A) Histograms on the left show phosphorylated STAT5 (p-STAT5) staining in day 22 cDC1. Control (blue) and Xcr1^Cre^STAT5a/b^fl/fl^ (red) iDC1 were re-stimulated with GM-CSF for 30 min, with unstimulated control cDC1 shown in gray. The graph on the right quantifies p-STAT5 mean fluorescence intensity (MFI). One of two independent experiments, each including 4-5 biological replicates per genotype, is shown. (B-D) Analysis of day 23 STAT5-deficient iDC1 cultures. Results are combined from two independent experiments, each including 4–5 biological replicates per genotype. (B) Representative contour plots showing CD45R/B220 and CD11c expression and percentages of cDC. (C, D) Quantification of relative (C) CD11c or (D) CD103 mean fluorescence intensity (MFI). (E) Histograms show BRD4 staining in day 22 cDC1 (left) or cDC2 (right). Control (blue) and Xcr1^Cre^Brd4^fl/fl^ (red) genotypes are compared, with unstained controls shown in gray. (F-H) Analysis of day 23 BRD4-deficient iDC1 cultures. Results are combined from two (CD103) or three (CD11c) independent experiments, each including 1–4 biological replicates per genotype. (F) Representative contour plots showing CD45R/B220 and CD11c expression and percentages of cDC. (G, H) Quantification of (G) relative CD11c or (H) CD103 mean fluorescence intensity (MFI). *** p<0.001, **** p<0.0001, NS = not statistically significant (unpaired two-tailed Student’s t-test).

**Table S1. Details of antibodies used in this study**

| **Antigen** | **Conjugate** | **Clone** | **Dilution** | **Supplier** | **Catalog #** |
| --- | --- | --- | --- | --- | --- |
| Brd4 | Unlabeled | EPR25425-23 | 1:50 | Abcam | ab289886 |
| Active caspase-3 | BV421 | C92-605 | 1:2560 | BD | 570786 |
| CD45R/B220 | PECy7 | RA3-6B2 | 1:320 | eBioscience | 25-0452-82 |
| CD45R/B220 | FITC | RA3-6B2 | 1:200 | BD | 553088 |
| CD3e | PE | 145-2C11 | 1:80 | BD | 553064 |
| CD3e | Biotin | 145-2C11 | 1:10 | BD | 553060 |
| CD8a | FITC | 53-6.7 | 1:200 | BD | 553031 |
| CD11b | eFluor450 | M1/70 | 1:160 | eBioscience | 48-0112-82 |
| CD11c | BV421 | N418 | 1:20 | Biolegend | 117330 |
| CD11c | Alexafluor594 | N418 | 1:400 | Biolegend | 117346 |
| CD4 | APC-eFluor780 | RM4-5 | 1:160 | eBioscience | 47-0042-82 |
| CD4 | APC | RM4-5 | 1:200 | eBioscience | 17-0042-83 |
| CD16/CD32 | 2.4G2 | Unlabeled | 1:10 | In house | N/A |
| CD19 | Biotin | eBio1D3 | 1:800 | eBioscience | 13-0193-85 |
| CD19 | APC-efluor780 | eBio1D3 | 1:160 | eBioscience | 47-0193-82 |
| CD24 | BV786 | M1/69 | 1:640 | BD | 744470 |
| CD40 | BV421 | 3/23 | 1:640 | BioLegend | 124641 |
| CD44 | APC | IM7 | 1:333 | eBioscience | 17-0441-83 |
| CD45 | BUV395 | I3/2.3 | 1:200 | BD | 567451 |
| CD45 | PECy7 | I3/2.3 | 1:1600 | BioLegend | 147704 |
| CD45.1 | FITC | A20 | 1:200 | Biolegend | 110706 |
| CD64 | Biotin | X54-5/7.1 | 1:400 | Biolegend | 139318 |
| CD80 | PECy7 | 16-10A1 | 1:640 | BioLegend | 104734 |
| CD86 | BV786 | GL1 | 1:800 | BD | 740877 |
| CD103 | APC | 2E7 | 1:640 | eBioscience | 17-1031-82 |
| CD115/CSF-1R | Alexafluor488 | AFS98 | 1:50 | BioLegend | 135512 |
| CD135 | APC | A2F10 | 1:10 | BioLegend | 135310 |
| CD172a | BUV805 | P84 | 1:320 | BD | 741997 |
| Clec9a | PE | 42D2 | 1:100 | eBioscience | 12-5975-82 |
| Clec9a | BUV395 | 7H11 | 1:10 | BD | 752673 |
| F4/80 | APC-eFluor780 | BM8 | 1:40 | eBioscience | 47-4801-82 |
| F4/80 | Biotin | BM8 | 1:200 | eBioscience | 13-4801-85 |
| Ly-6G/Ly-6C | APC-eFluor780 | RB6-8C5 | 1:40 | eBioscience | 47-5931-82 |
| H2Kb | FITC | AF6-88.5 | 1:200 | BD | 553569 |
| I-A/I-E | PE | M5/114.15.2 | 1:2560 | Biolegend | 107607 |
| I-A/I-E | Alexafluor647 | M5/114.15.2 | 1:3200 | BioLegend | 107618 |
| I-A/I-E | BUV805 | M5/114.15.2 | 1:200 | BD | 748844 |
| IgG1k Isotype | Alexafluor489 | MOPC-21 | 1:2 | BD | 557782 |
| Ly-6G | Biotin | 1A8 | 1:100 | Biolegend | 127604 |
| NK1.1 | Biotin | PK136 | 1:200 | BD | 553163 |
| NK1.1 | APC-efluor780 | PK136 | 1:50 | eBioscience | 47-5941-82 |
| CD170/SiglecF | Biotin | S17007L | 1:100 | Biolegend | 155512 |
| STAT5 | Alexafluor488 | 47/Stat5 (pY694) | 1:2 | BD | 612598 |
| TCRb | Biotin | H57-597 | 1:200 | BD | 553169 |
| TCRb | Alexafluor594 | H57-597 | 1:400 | BioLegend | 109238 |
| Ter119 | APC-efluor780 | TER-119 | 1:40 | eBioscience | 47-5921-82 |
| Ter119 | Biotin | TER-119 | 1:400 | eBioscience | 13-5921-82 |
| Va2 TCR | PE | B20.1 | 1:320 | Invitrogen | 12-5812-82 |
| Xcr1 | BV510 | ZET | 1:80 | Biolegend | 148218 |
